# Treeline ecotone habitat loss and upward shift is driven by land abandonment rather than climate change in the Italian Alps

**DOI:** 10.64898/2026.09.30.755293

**Authors:** Carrieri Erik, Ha Trang Nguyen, Meloni Fabio, Anselmetto Nicolò, Morresi Donato, Garbarino Matteo

## Abstract

1. Plant populations at high elevation persist near their physiological limits, and treeline position is one of the clearest expressions of this constraint. In many mountain regions, however, this limit reflects historical land use as much as climate. Treeline shift is most often tracked as the displacement of a boundary line (1D) overlooking structural changes only visible when the ecotone is analysed as a surface (2D).
2. We reconstructed treeline ecotone dynamics at nine sites in the Italian Alps using aerial imagery from 1954 and 2017. At each site we detected the timberline and tree species line, quantified elevational displacement, and measured ecotonal surface change, relating them to climatic, land-use, and topographic predictors.
3. Both boundaries advanced upslope at all nine sites, with timberlines shifting faster (1.6 m yr ¹) than tree species lines (1.1 m yr ¹). This decoupled velocity caused the ecotonal surface to contract by ∼17%. Land-use variables consistently explained more variance than climatic ones.
4. Land-use legacy and abandonment have a major role in shaping treeline dynamics. Boundary displacement reveals vegetation advance at both margins, while the surface view reveals concurrent narrowing of the habitat available for tree establishment. Both perspectives are needed to fully characterize how plant populations respond to shifting limiting factors at the treeline ecotone.

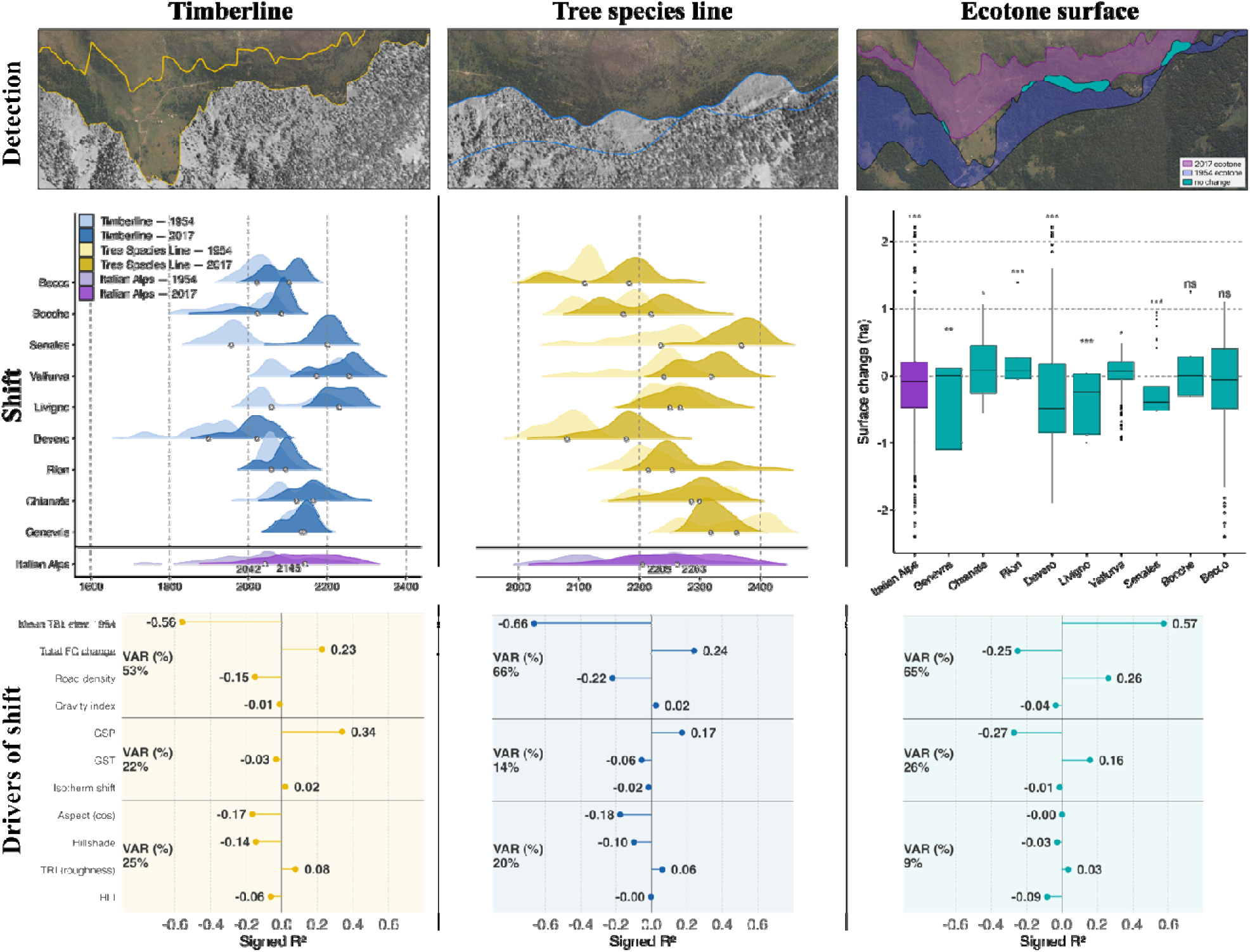

## 1 | Introduction

Plant populations at high elevation persist at the edge of their physiological tolerance, and the position of the treeline is one of the clearest expressions of this limit. Forest upper elevational boundaries are shifting worldwide as a result of global environmental changes. As these changes affect alpine plant communities and species composition (Garbarino et al., 2022; Tasser et al., 2025; Vanneste et al., 2026), understanding the role of the driving factors on such dynamics is a core challenge in ecology. Increasing growing season temperature driven by climate change is broadly recognized as one of the main drivers of treeline position (Hansson et al.,2023, Schwörer et al., 2014, Xie et al., 2024). As a consequence of the warming of mountain regions (Beniston, 1997; Chan et al., 2024), it is expected for treelines to respond by advancing upslope into formerly inhospitable areas; however, many treelines at the global level are failing to track the isotherms shift (Xie et al., 2024).

The asynchronous response to global warming reflects the multifaceted nature of treeline ecotones, as other factors like land-use history and topography locally mediate broad-scale climate patterns (Garbarino et al., 2023; Carrieri et al., under revision). The effect of topography on high-altitude forest dynamics is well documented (Beniston, 2003; Resler, 2006; Tourville et al., 2023), as it can influence wind exposure (Beloiu et al., 2022), radiation stress (McIntire et al., 2016), snow cover duration (Batllori et al., 2009), shelter provision (Batllori et al., 2009; Holtmeier and Broll, 2007), and fine-scale temperatures (Hamid et al., 2023; Marquis et al., 2021). Concurrently, terrain features serve as a proxy of past human pressure. In the European Alps, pristine treelines are often restricted to steep, rough, and rocky slopes where alpine farming and forest pasturing were historically less diffuse (Ameztegui et al., 2016; Gehrig-Fasel et al., 2007; Tasser et al., 2007).

Land-use practices have been acknowledged as primary drivers of treeline ecotone dynamics in many alpine environments (Ameztegui et al., 2016; Delpouve et al., 2025; Tasser et al., 2007). Recent studies highlight how land-use legacies can shape treeline elevation (Frei et al., 2023; Liang et al., 2026) and that the relaxation of intense human activities (e.g., intensive grazing) resulted in faster shifts than those recorded in less disturbed areas (Ameztegui et al., 2016; Delpouve et al., 2025). Although rising temperatures provide the broad-scale climatic potential for upslope advance, the realized treeline movement may also depend on where and when land use has recently changed, making it difficult to attribute observed shifts unambiguously to either driver.

In the European Alps, the boundary between the continuous closed forest and the alpine grassland, which we here refer to as “timberline” (canopy cover >80%), and the “tree species line”, the line connecting the uppermost detectable (crown size > 4 m2, 2×2 pixels) trees regardless of species, have different histories and dynamics. In many European Alpine regions, land-use history has lowered the timberline through millennia of grazing, mowing and woodcutting (Kyriazopoulos et al., 2017; Lasanta et al., 2017; Wielgolaski et al., 2017). Since the mid-19th century, declining pastoralism and rural abandonment in the European Alps have triggered widespread forest regrowth at high altitudes (Anselmetto et al., 2024, submitted manuscript; Mather, 1992; Tasser et al., 2007), and the combined effect of land abandonment and recorded temperature increases since the 20th century has further boosted tree encroachment in high-altitude environments (Gehrig-Fasel et al., 2007; Piccinelli et al., 2020). Disentangling the relative contribution of climate warming and land-use change to treeline dynamics remains a major challenge, particularly for anthropogenic treeline ecotones that dominate the European Alps, where the two drivers often act simultaneously (Gehrig-Fasel et al., 2007).

Research on treeline range shifts mainly focuses on the shift of the “treeline” intended as a one-dimensional boundary line (Garbarino et al., 2023; He et al., 2023; Xie et al., 2024), measuring its displacement in terms of elevational change of either the tree species line or the timberline (Hansson et al., 2021; Harsch et al., 2009; Xie et al., 2024), while overall ecotonal surface changes are often overlooked. However, elevational shifts are complex phenomena, and the spatio-temporal dynamics of the upper and lower boundaries may differ (Birre et al., 2023; Rumpf et al., 2018). Multi-dimensional perspectives can capture vegetation spatial patterns reflecting structural changes, underlying processes, and environmental heterogeneity that elevation shifts of linear boundaries (1D) alone fail to capture (Bader et al., 2021; Birre et al., 2023) and can therefore be used to better understand ecotonal shifts (Hooshyarkhah et al., 2025) and their consequences. As trees at the tree species line —upper boundary of the treeline ecotone— grow closer to their thermal niche, a relaxation of anthropogenic pressure at the timberline may favor a faster upshift of the lower boundary, which is more related to disturbance regimes than to climatic limitations, resulting into a measurable reduction of the ecotonal surface area over time.

Here we present a comprehensive assessment of the spatio-temporal dynamics of treelines—both as a line and as an ecotone—and their drivers in the Italian Alps over six decades. We explore recent ecotonal changes in terms of timberline and tree species line position, and ecotonal surface area, through a comparison of diachronic aerial photography from 1954 and 2017 across nine Alpine sites. We examine the pattern and extent of treeline change to investigate the role of historical land use, climate warming, and topography on the observed temporal dynamics of realized treeline elevation. We hypothesised that shifts interesting the treeline ecotone elevation from the 1950s are: (1) still decoupled from the global warming signal in the Alps, (2) mostly modulated by land-use legacy and post-abandonment natural forest expansion at the landscape scale, rather than with the recent warming signal alone, and (4) characterised by different velocities between the two boundaries, with the timberline advancing faster than the tree species line, leading to the loss of ecotone surface over time.

## 2 | Materials and Methods

### 2.1 | Study Area

The combination of strong climatic and topographic heterogeneity with the long history of human land-use makes the Italian Alps ideal to investigate treeline ecotone dynamics (Anselmetto et al., 2024; Bätzing et al., 1996). Nine landscapes were selected along an East–West gradient spanning the Western, Central and Eastern sectors of the Italian Alps, capturing broad climatic, topographic, edaphic, vegetation, and land-use pressure heterogeneity (Fig. 1). Therefore, the study sites offer a gradient of conditions suited to disentangling the relative role of drivers of change. Given the marked orographic variability among sites, the extent of the treeline ecotone differs across landscapes (Table S1), with a total investigated area of 106 km². Sites span an elevation range from 2050 m a.s.l. (Genevris) to 3010 m a.s.l. (Chianale), capturing the transition zone from closed subalpine forest to the upper tree limit — the belt widely documented as the treeline ecotone in the European Alps (Hättenschwiler and Körner, 1995; Paulsen et al., 2000). The study sites encompass high variability in terrain features including aspect, slope, and roughness, and exhibit contrasting precipitation regimes (< 1000 mm yr ¹ at Chianale, Senales, and Valfurva; > 1100 mm yr ¹ at the remaining sites). Dominant species are subalpine conifers, primarily European larch (*Larix decidua* Mill.), Swiss stone pine (*Pinus cembra* L.), Norway spruce (*Picea abies* L.), and Scots pine (*Pinus sylvestris* L.). Dominant species and site features were validated through field campaigns conducted as part of a previous study (Carrieri et al., 2025). Further details on the source and post-processing of the 1-m resolution 1954 grayscale aerial photographs and 2017 RGB orthophotos are provided in Supplementary Material A.

**Figure 1.**
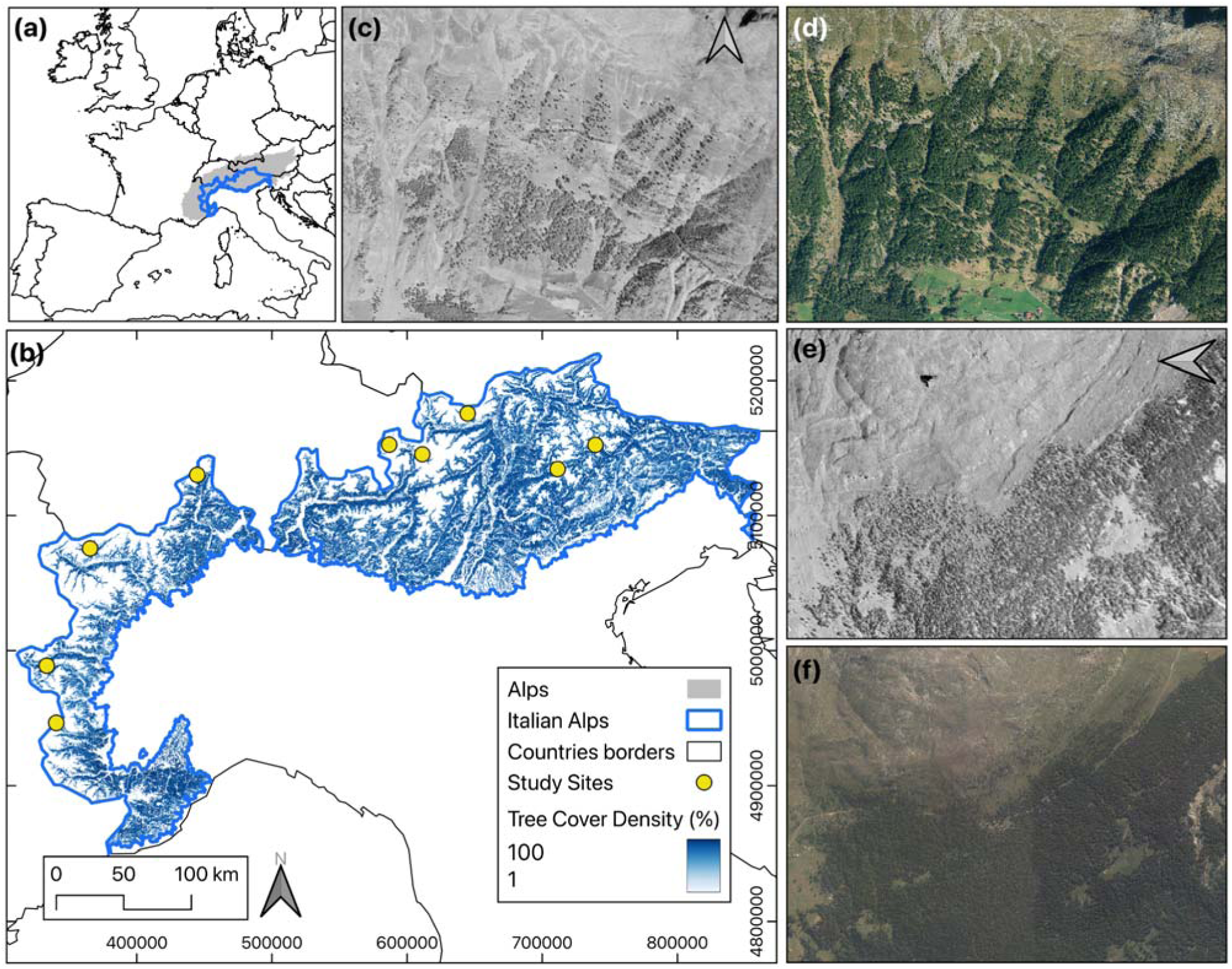
| Locations of the study sites. Map of the broader study area within the Alpine Region (a); locations of the nine study sites displayed as circles superimposed on the tree cover density in the Italian Alps with a white-to-blue density gradient (b); Senales landscape in 1954 (c) and in 2017 (d); and Devero landscape in 1954 (e) and 2017 (f). Both case studies exemplify the broader trend of afforestation observed throughout the Alpine region.

### 2.2 | Treeline detection

To delineate the timberline and tree species line, we produced a binary tree/non-tree cover layer for each time period using a UNet++ convolutional neural network (Fig. 2; Zhou et al., 2018). Two separate models were trained — one for each time period — using manually annotated tiles (96 RGB, 91 grayscale) stratified to represent variability in crown morphology, illumination, and background texture (Fig. S1). The encoder was an SE-ResNeXt101 backbone pretrained on ImageNet. Models were optimized using the Lovász loss function with the Adam optimizer (learning rate 1×10). Performance was evaluated with IoU, F1 score, and Cohen’s Kappa. Model outputs were manually post-processed to remove residual false positives (isolated patches on bare ground or rocky slopes) and to fill shadow-induced classification gaps via focal statistics (Fig. S2-3; Table S2). Full details of the training protocol, annotation criteria, deep learning model, and post-processing workflow are provided in Supplementary Material B.

**Figure 2.**
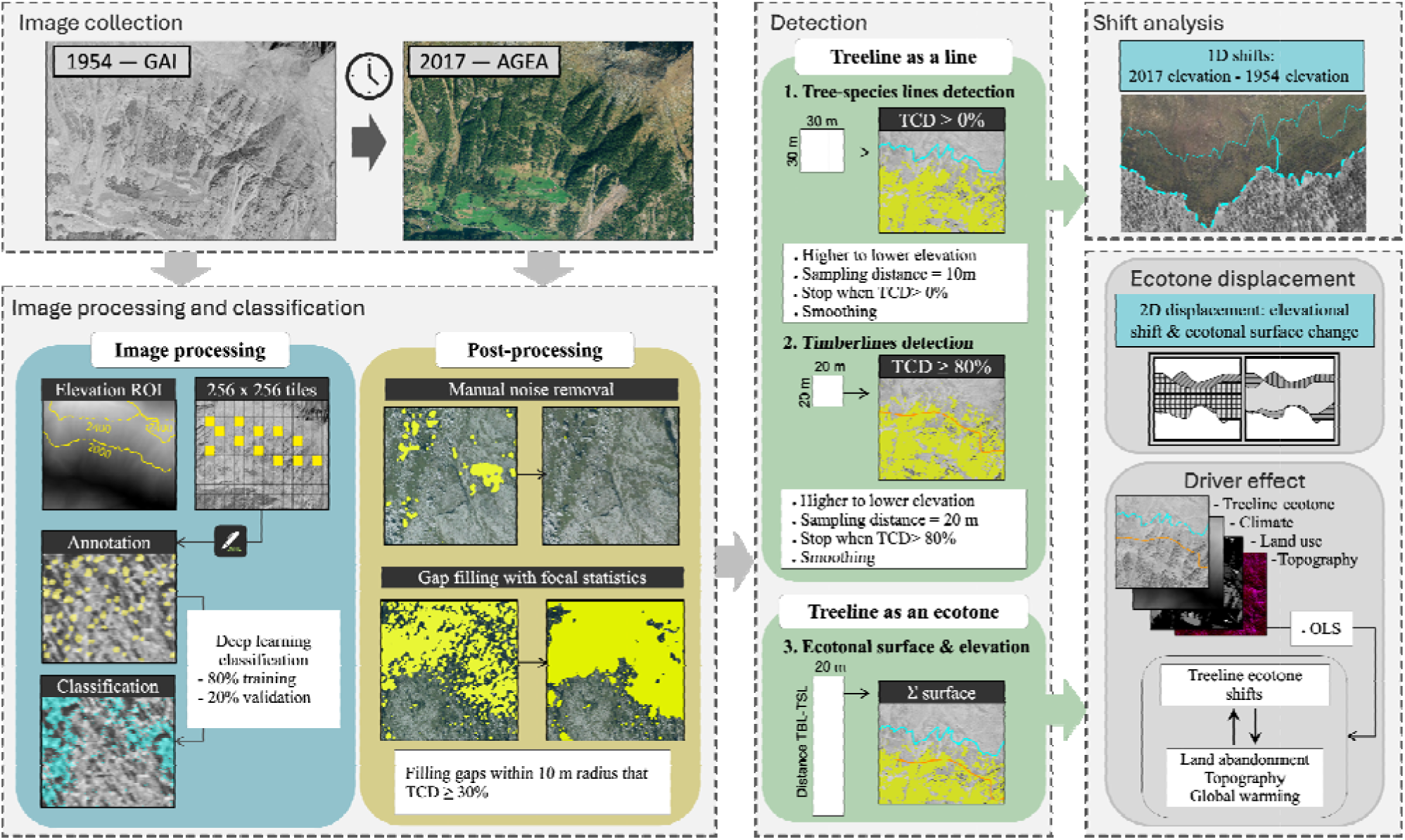
| Methodological workflow of the study to delineate the treeline ecotone and analyse its elevational shifts. Image collection: compilation of 1954 grayscale aerial photographs (GAI flight) and 2017 RGB orthophotos (AGEA). Image processing and classification: definition of a region of interest (ROI), tiling, manual annotation, deep-learning classification, and post-processing (noise removal, gap filling) based on tree cover density (TCD) thresholds. Detection: identification of the tree species line and timberline detected as 1D lines at increasing TCD thresholds and the ecotonal surface, computed as the surface between the two lines. Shift analysis: quantification of 1D elevational (2017–1954) and 2D ecotone shifts between the two time steps. Driver effect: relationship between treeline ecotone shift and drivers of change via ordinary least squares (OLS) regression.

### 2.3 | Treeline shifts measurement

#### 2.3.1 | Treeline as a line

Elevational shifts of the timberline and tree species line were quantified using a transect-based moving-window approach similar to the one proposed by Gehrig-Fasel et al. (2007). A set of transects spaced 10 m apart were generated across each region of interest (ROI), and a 30 × 30 m moving window was scanned downslope along each transect in 10-m steps. At each step, the proportion of tree pixels from the modelled binary tree cover was evaluated within the window. The tree species line was defined as the position of the highest elevation tree occurrence per transect, while the timberline was defined as the upper limit of canopy cover exceeding 80% within the window. Elevational shift per each transect was computed as the difference between the 2017 and 1954 DEM-derived elevations, with positive values indicating upslope displacement (Fig. 3a–b). Site-level overall tree species line and timberline elevational shifts were produced by computing median per-transect shifts for each line.

**Figure 3.**
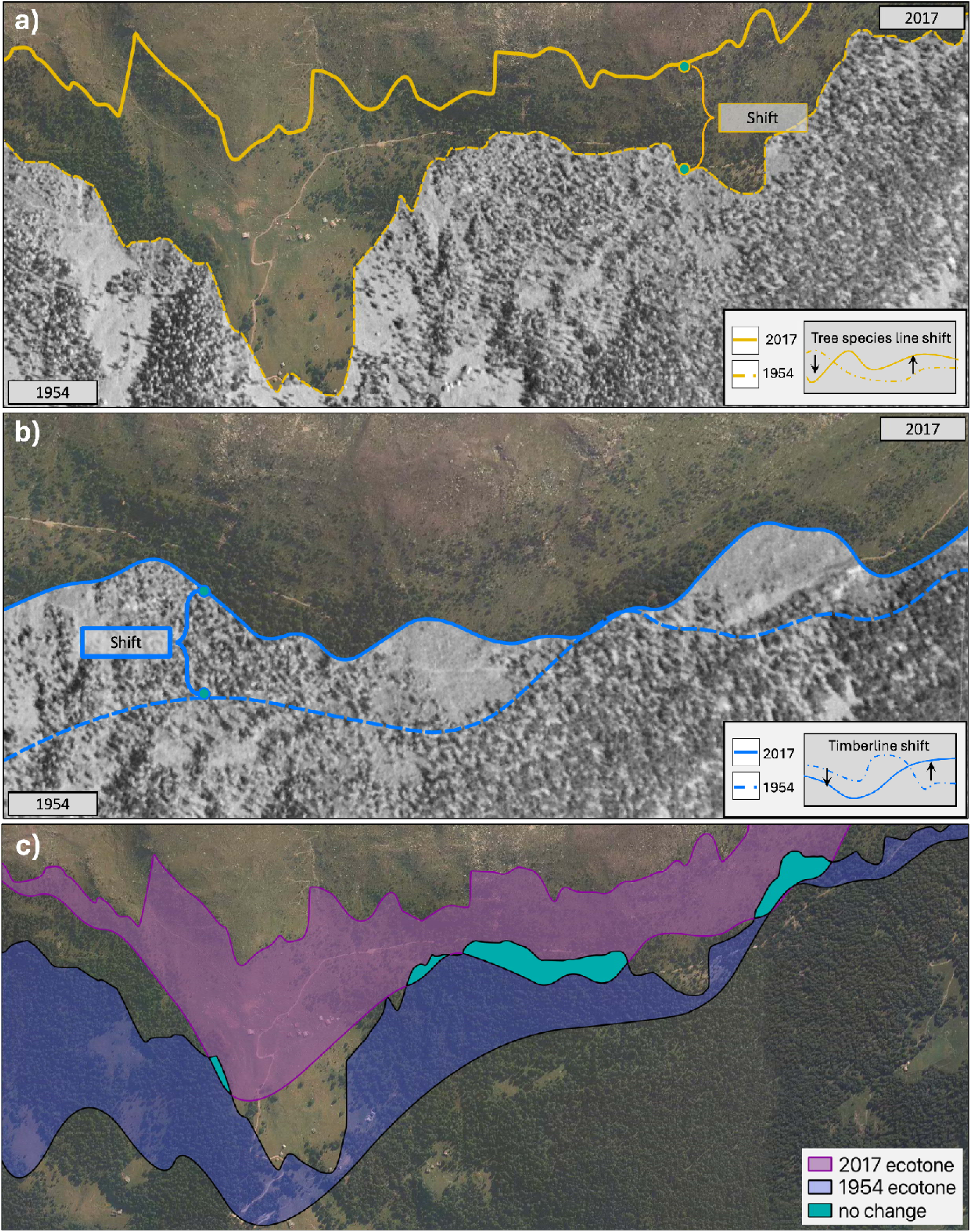
| Illustration of the three ecotonal change metrics measured in this study. Panel (a) shows the upslope displacement of the tree species line. The cyan solid line marks the 2017 boundary and the cyan dashed line marks the 1954 boundary. Panel (b) shows the upslope displacement of the timberline, with the 2017 boundary in solid orange and the 1954 boundary in dashed orange. Panel (c) shows the ecotone surface change. The gold polygon represents the ecotone extent in 1954, and the blue polygon represents the ecotone extent in 2017. The grey area indicates the portion of the ecotonal surface present in both time periods that has not changed in extent. Areas covered exclusively by the gold polygon represent ecotone surface lost since 1954, while areas covered exclusively by the blue polygon represent newly gained ecotone surface.

#### 2.3.2 | Treeline as a bidimensional space

Ecotonal surface change and elevational shift were measured leveraging the same linear transects used to measure the boundaries shifts. For each transect, an ecotone polygon was constructed by identifying the intersection points of the transect line with the timberline and tree species line layers for each time period. The segment connecting these two intersection points was buffered by 10 m on each side using flat end caps, yielding a rectangular strip representing the ecotone width along that transect—equivalent to a 20-m wide field transect spanning from the closed forest boundary (i.e., timberline) to the uppermost detectable tree (i.e., tree species line). Ecotone surface change was computed as the difference in polygon area between 2017 and 1954, with positive values indicating expansion (Fig. 3c). Ecotone elevational shift was computed as the difference between the 2017 and 1954 median DEM elevation within each polygon. Both metrics were summarized at site level as the median per-transect value to ensure comparability across sites with different transect counts.

### 2.4 | Driving factors of treeline ecotonal change

#### 2.4.1 | Predictors

To explore the drivers of ecotonal shifts in elevation and surface, elevational and surface changes in the treeline ecotone were related to climatic, land-use, and topographic predictors through a series of complementary statistical models.

Three categories of predictor were compiled at the site level (Table 1). Climatic predictors comprised growing-season temperature (GST) and growing-season precipitation (GSP), extracted from CHELSA bioclimatic rasters (Karger et al., 2017), and the annual upward displacement of temperature isotherms (Chan et al., 2024) as an indicator of recent thermal shift. Highly collinear CHELSA variables (growing season length (GSL), growing degree days (GDD), snow cover duration (SCD); pairwise |r| > 0.85) were excluded following an initial correlation screening (Fig. S4). Land-use predictors included the total proportional forest cover change between 1954 and 2017 (derived from Anselmetto et al., submitted manuscript), the mean elevation of the 1954 timberline as a proxy for historical land-use legacy, and road density (m km ²) within a 5 km site buffer derived from OpenStreetMap. Among land-use predictors we included a population gravity index (Polyakov et al., 2008) to explore the combined effects of size and proximity of the population (Supplementary Material C). Topographic predictors were derived from the 10 m DEM and included the topographic ruggedness index (TRI), hillshade, the cosine component of aspect, and the heat load index (HLI; McCune and Keon, 2002).

**Table 1.**
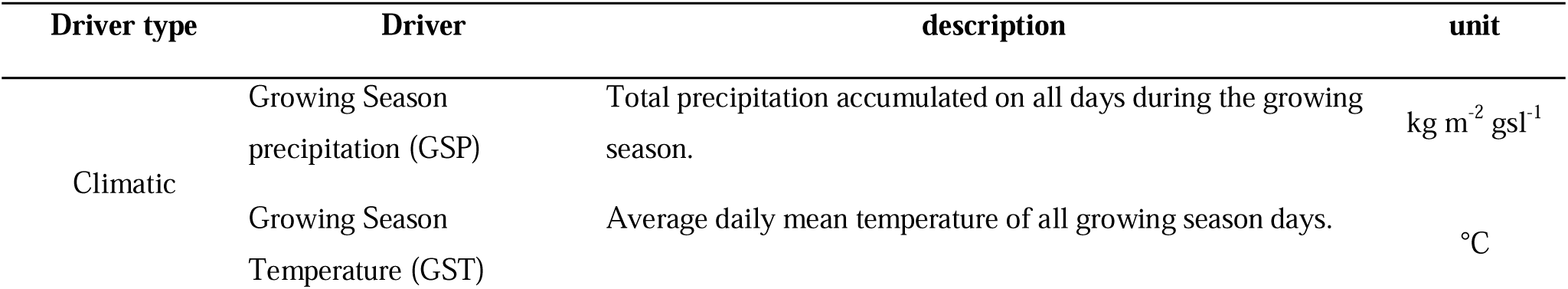
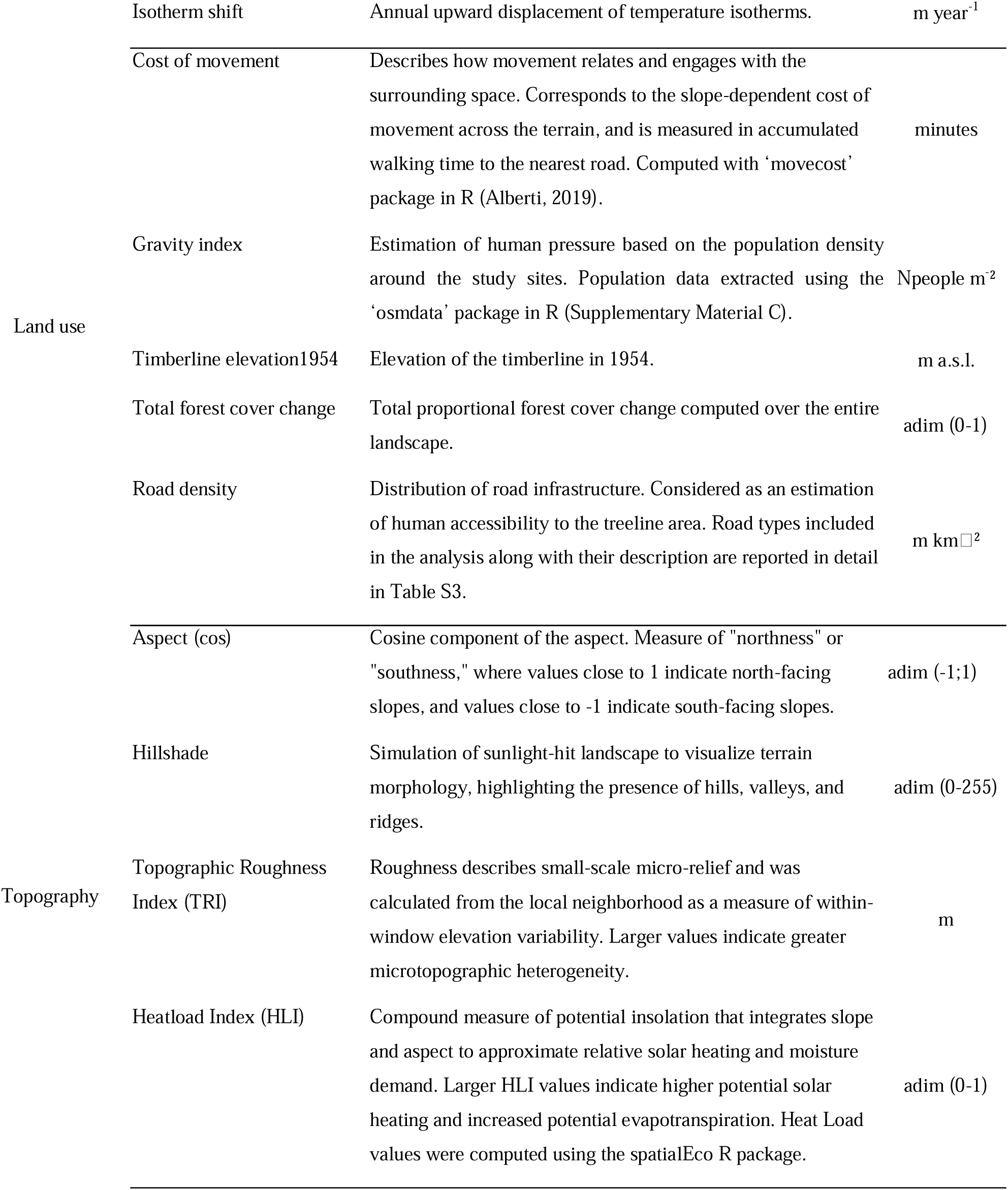
| Suite of climatic, land use, and topographic predictors of timberline and tree species line shift and ecotonal surface change grouped by driver type. For the growing season precipitation and growing season temperature, the growing season length was defined as the number of days between the first and last occurrence of growing conditions, where the growing conditions is a mean daily air temperature above 0.9°C according to the climate-based model “TREELIM” (Paulsen &Körner, 2014).

#### 2.4.2 | Statistical analysis

All predictors were normalized prior to modelling to allow direct comparison of effect sizes. For each of the three response variables — timberline elevational shift, tree species line elevational shift, and ecotone surface change — simple ordinary least squares (OLS) regressions were fitted one predictor at a time. Fitting multivariate models or non-linear approaches (e.g. GAMs) was not feasible given the site-level sample size (n = 9). Effect direction was preserved by reporting signed R², computed as sign(β) × R². Spearman rank correlations (ρ) were computed alongside each regression as a non parametric robustness check, against the influence of outlying sites. Residual normality and homoscedasticity were assessed using the Shapiro-Wilk and Breusch-Pagan tests respectively (Table S4). All analyses were performed in R v4.2.1.

## 3 | Results

### 3.1 | Timberline, tree species line and treeline ecotonal shifts

Both timberline and tree species line have shifted upward over the last 60 years, with elevation shift rates of 1.6 m year^-1^ and 1.15 m year^-1^, respectively (Fig. 4-5; Table 2). In two thirds of the sites, the lower boundary —timberline— shifted faster than the upper one —tree species line. As a result, the ecotone surface tends to be 17% narrower nowadays than in the 1950s. Overall, 96% of all the transects showed an upslope shift, and 56% recorded an ecotonal surface contraction (Fig. 6; Table 2). The strongest boundary shifts were observed at Senales, where the median timberlines advance is 250 m, and the median tree species line shift was ∼150 m, while Genevris showed the lowest shift in both boundaries (<10 m). At Devero, both boundaries advanced largely (123 m the timberline, 89 m the tree species line). More moderate advances were observed at Chianale, Rion, Livigno, Valfurva, Becco, and Bocche. In Genevris, Rion, and Valfurva the tree species line advanced faster than the timberline Overall, the total ecotonal surface area of the entire region contracted by 149 ha (17%), with a rate of 2.38 ha year^-1^ (–0.27% year^-1^). Five out of nine sites (Devero, Genevris, Livigno, Becco and Senales) showed a reduction in the total ecotonal area (Fig. 6a; Table 2), with Senales, Livigno, and Devero exhibiting the highest surface loss (∼0.5 ha year^-1^). Surface gain was recorded in Chianale and Rion (∼0.1 ha year^-1^), while Valfurva and Bocche showed stable ecotonal surface.

**Figure 4.**
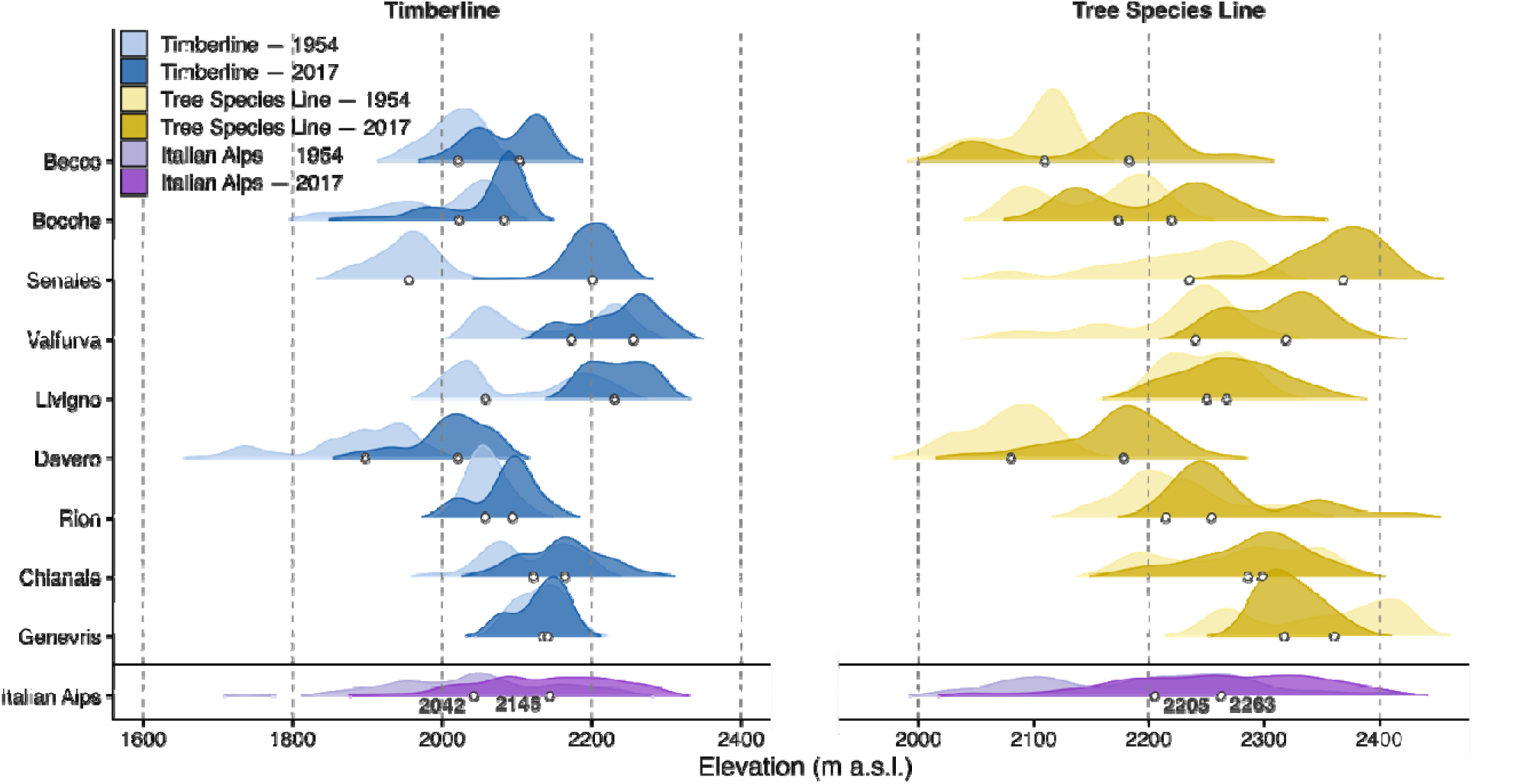
| Kernel density distributions of the absolute elevation of the timberline (blue, left panel) and tree species line (gold, right panel) in 1954 (light shading) and 2017 (dark shading) across the nine study sites and pooled across the Italian Alps (purple, bottom row). Each distribution summarises observations from all transects within a site, where each observation corresponds to the elevation at which the timberline or tree species line intersects a single transect. Broader and taller portions of a distribution indicate a higher density of transect observations at that elevation. White dots denote the median elevation of each distribution.

**Figure 5.**
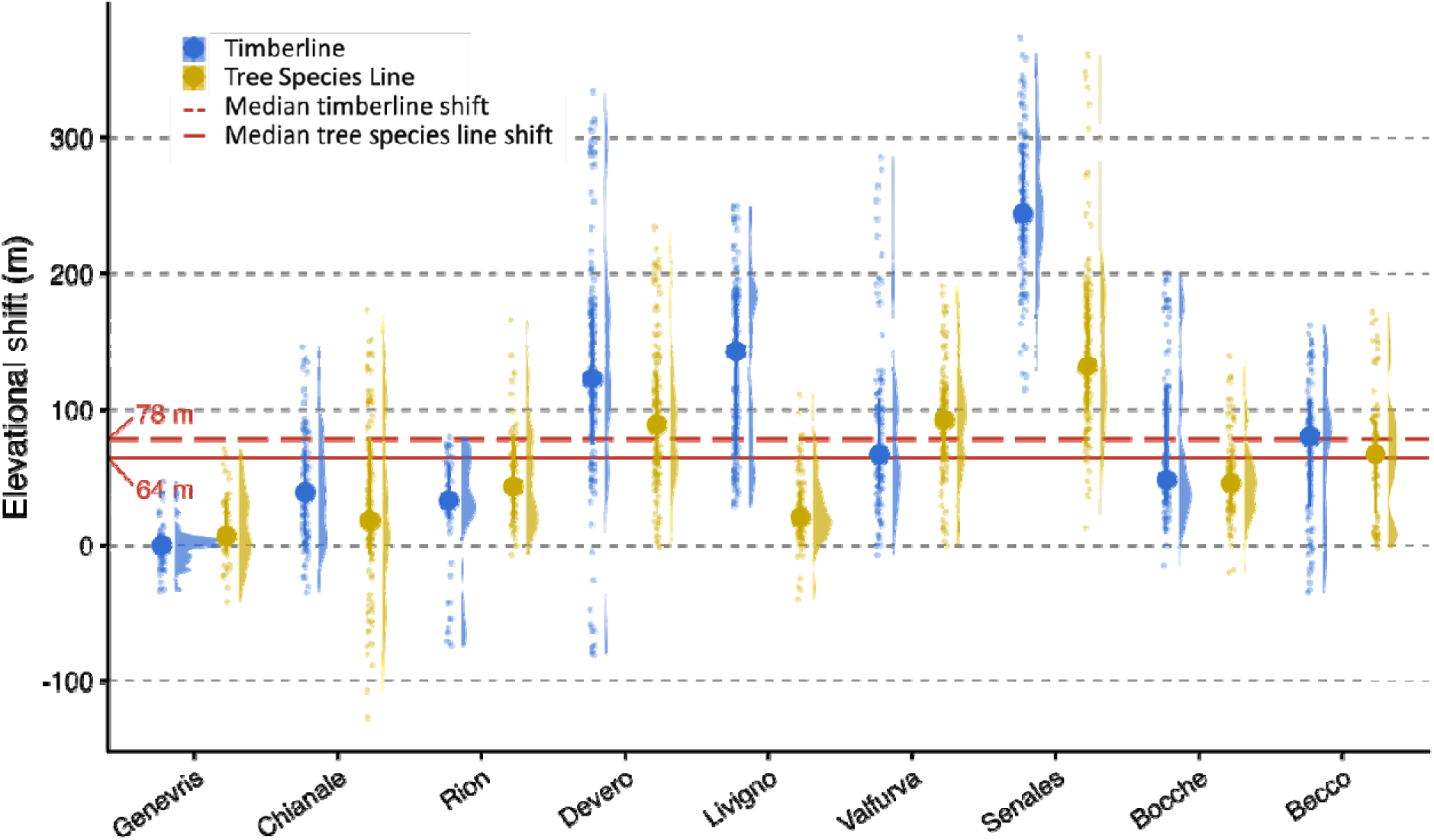
| Per-transect elevational shift (m) of the timberline (blue) and tree species line (gold) between 1954 and 2017 across the nine study sites. Each point represents the elevational shift measured at a single transect computed as the difference between the 2017 and 1954 elevation. Half-violins to the right of each point cloud shows the kernel density distribution of transect-level shifts, with taller portions indicating a higher density of observations around that value. The central dot on the point cloud denotes the site median. Horizontal red lines indicate the median elevational shift pooled across all transects and sites in the Italian Alps for the timberline (dashed) and for the tree species line (solid).

**Figure 6.**
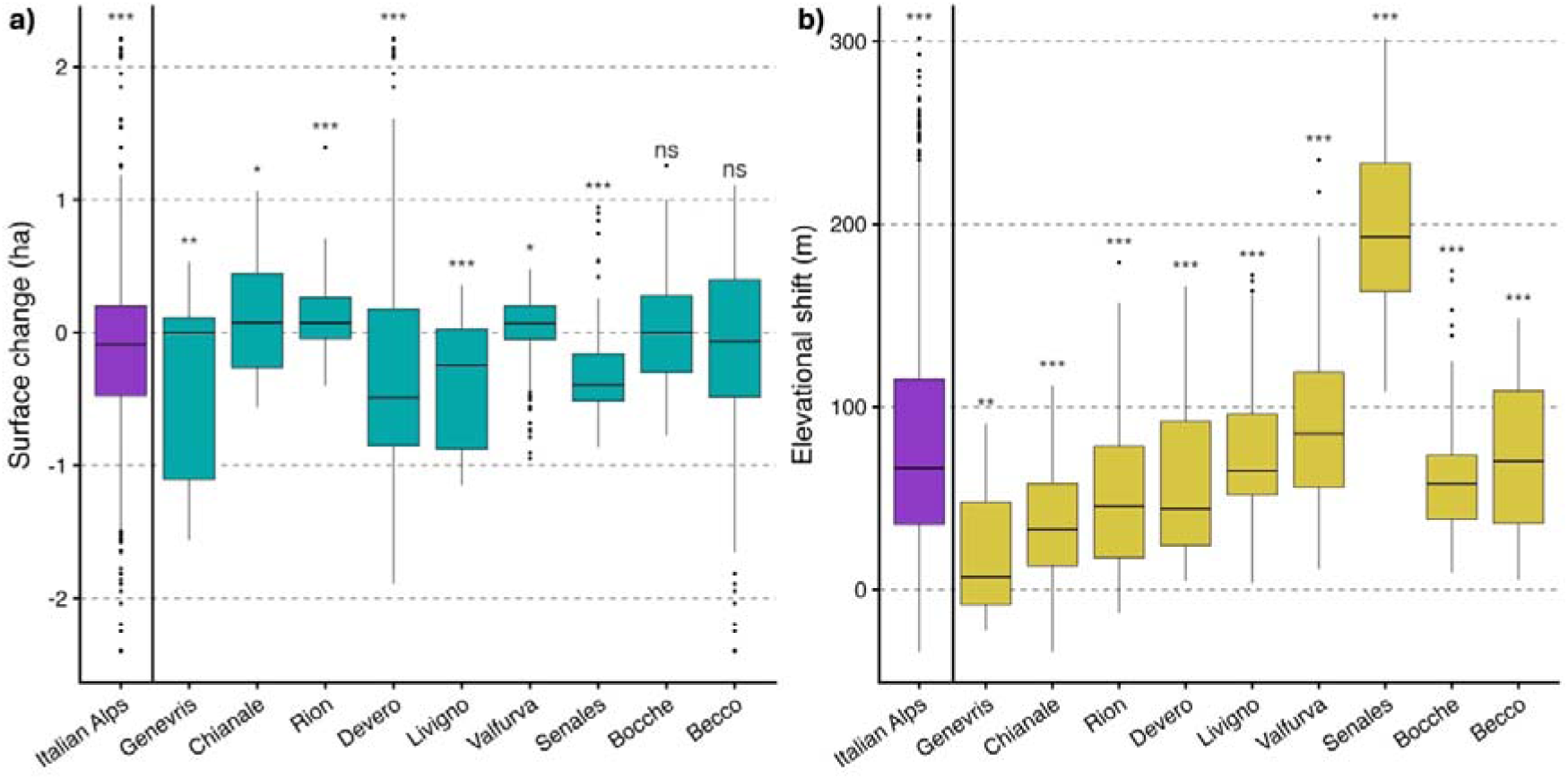
| Per-transect ecotone surface change (a) and ecotone elevational shift (b) between 1954 and 2017, shown for each of the nine study sites and pooled across the Italian Alps (purple). Each observation corresponds to a single transect shift. The horizontal black line within each box indicates the median. Box edges represent the interquartile range (IQR: 25th-75th percentiles). Whiskers extend to the largest and smallest values within 1.5 × IQR from the box edges. Stars above each box indicate the result of a Wilcoxon signed-rank test against the null hypothesis of no change. \**p* < 0.05; \*\**p* < 0.01; \*\*\**p* < 0.001; ns = not significant.

**Table 2.**
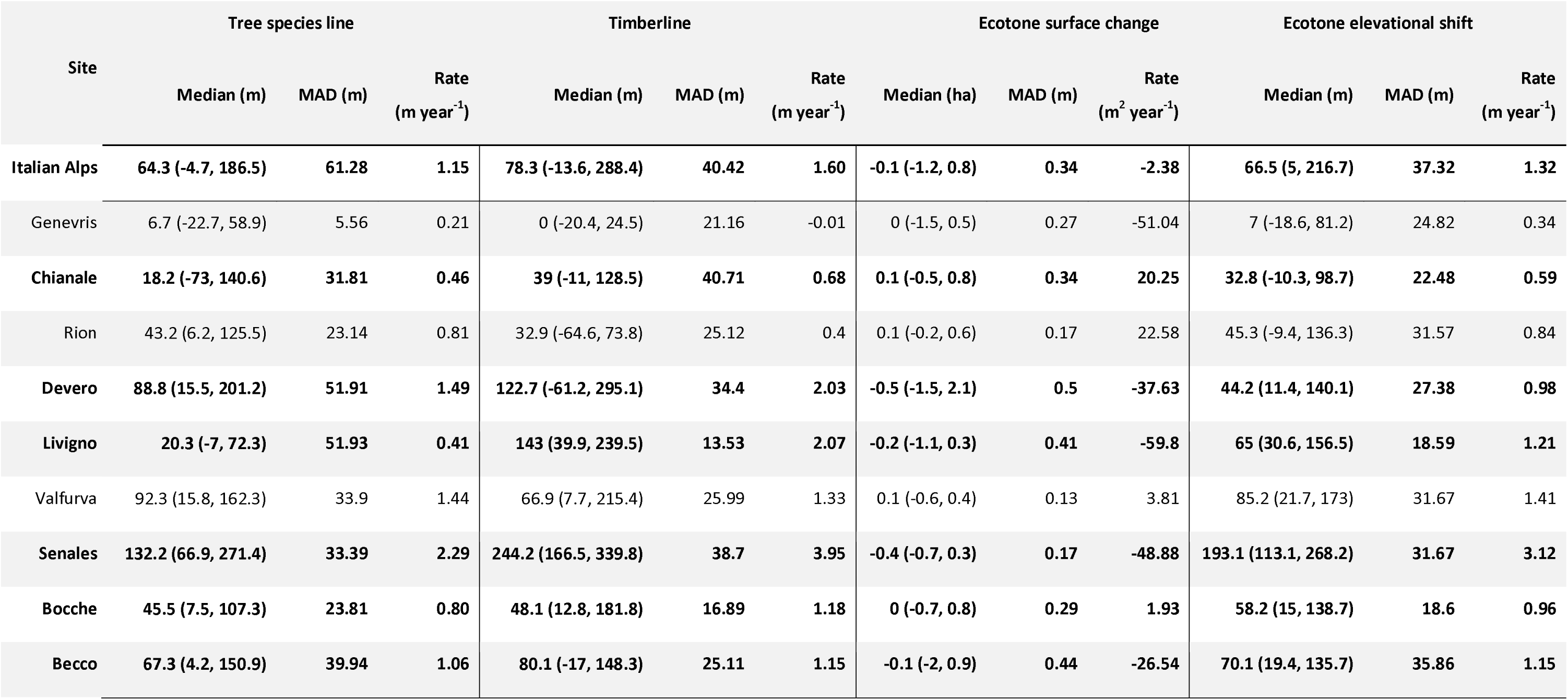
| Tree species line and timberline shift and ecotonal surface change and elevational shift. For each response variable we report medians (with 5th and 95th percentile in brackets), mean absolute deviation (MAD), and rate of the shift across all study sites. Sites with timberline shift greater than tree species line shift are highlighted in bold.

In terms of elevational displacement (Fig. 6b), all the ecotones shifted upward across all sites, although with varying magnitude (Table 2). Senales showed the strongest elevational gain (mean ∼3 m year^-1^), followed by Valfurva (∼1.5 m year^-1^). Becco, Bocche, Livigno, Rion and Devero exhibited moderate upward shifts (∼1 m year^-1^), whereas Chianale and Genevris recorded relatively small migration (∼ 0.5 m year^-1^).

### 3.2 | Driving factors of ecotonal changes

Across all three response variables, land-use ones retained the greatest explanatory power (53%, 66%, and 65% of the total variance of the tree species line, timberline, and ecotone dynamics, respectively), with the baseline elevation of the 1954 timberline emerging as the dominant predictor of both elevational and surface dynamics of the treeline ecotone (Table S5). Timberline and tree species line shifts were found to be consistently influenced by all the variables in the same manner in terms of magnitude and direction of the association (Fig. S5-S6). Among land-use variables, the mean elevation of the 1954 timberline was the strongest predictor of tree species line shift (R² = 0.56; ρ = −0.57). Total forest cover (FC) change explained a further 23% of the variance (R² = 0.23; ρ = 0.37), with landscapes undergoing greater forest expansion showing larger advances. Road density ranked third (R² = 0.15; ρ = −0.33), suggesting greater tree species line advancement in more remote sites. Climate explained more variance of the tree species line shifts (22%) than of the timberline ones (14%). Among climatic predictors, GSP was the only significant variable (R² = 0.34; ρ = 0.70). GST and isotherm shift explained little variance (R² = 0.03 and 0.02, respectively). Among topographic variables, aspect (cosine) explained the most variance (R² = 0.17; ρ = −0.47), followed by hillshade (R² = 0.14; ρ = 0.25), while TRI and HLI both had R² values < 0.15.

For timberline shifts, the mean elevation of the 1954 timberline again emerged as the dominant predictor (R² = 0.66; ρ = −0.62). Total FC change and road density showed moderate associations (R² = 0.24 and 0.22; ρ = 0.48 and −0.35, respectively). GSP remained the strongest climatic predictor (R² = 0.17; ρ = 0.63), whereas GST and isotherm shift each explained <6% of the variance. Aspect (cosine) was again the strongest topographic predictor (R² = 0.18; ρ = −0.40), while hillshade, TRI, and HLI all had R² values < 0.10.

The pattern of driver importance for ecotonal surface change diverged from the boundaries shifts responses (Fig. S7). Mean elevation of the 1954 timberline remained the strongest predictor (R² = 0.57; ρ = 0.67), but with a positive relationship, indicating that higher-elevation treeline ecotones gained surface area. Land-use variables showed weaker, reversed associations relative to elevational shifts, with total FC change (R² = 0.25; ρ = −0.30) and road density (R² = 0.26; ρ = 0.23) showing opposite effects. GSP was again the strongest climatic predictor and second overall (R² = 0.27; ρ = −0.58), with wetter sites tending toward greater ecotone contraction. GST showed a moderate positive association (R² = 0.16; ρ = 0.25), whereas isotherm shift explained virtually no variance (R² = 0.01). Topographic variables were consistently weak predictors, with all R² values < 0.10.

## 4 | Discussion

Our results indicate that, over the past 60 years, Italian Alpine treelines shifted upward at both their lower and upper boundaries, and that treeline ecotone surface has substantially narrowed. This shrinking of the ecotones —17% narrower today than in the 1950s— is linked to a decoupled velocity of their boundaries driven by anisotropic response of vegetation to the removal of the same limiting factor (i.e., anthropogenic disturbance). As a result, timberlines —the lower boundaries— have advanced faster than tree species lines (1.6 vs 1.1 m year ¹). Historical aerial photographs coupled with computer vision techniques allowed us to unveil the legacy of historical human pressure on vegetation as the primary explanatory factor of treeline dynamics in this region.

### 4.1 | Upward shift of treeline and ecotonal shrinkage

Although upward treeline shifts are widely documented across the Alps (Frei et al., 2023; Gehrig-Fasel et al., 2007; Körner and Hiltbrunner, 2024), this study is the first to quantify the different rates of timberline and tree species line advance and relating this asymmetry with the contraction of the ecotonal surface (Fig. 5). We argue that overlooking the bi-dimensional component of the treeline ecotone risks missing an important dimension of treeline change: a narrowing ecotone implies a compression of the transition zone between closed forest and alpine grassland, with potential consequences on its structural complexity, habitat diversity, and buffering capacity of this zone against further environmental change (Dirnböck et al., 2003).

The elevational shift rates recorded in this study agree with recent estimates of Alpine treeline advance of approximately 1 m year ¹ (Delpouve et al., 2025; Frei et al., 2023; Menegaldo et al., 2025). Although differences in the rates of timberline and tree species line movement have rarely been investigated (but see Hooshyarkhah et al., 2025), the asymmetric boundary response and resulting ecotone contraction are consistent with previous observations (Delpouve et al., 2025; Dirnböck et al., 2003; Rumpf et al., 2018).

Our findings align with a growing body of research adopting multi-dimensional perspectives on treeline dynamics. Feuillet et al. (2020) and Delpouve et al. (2025) similarly moved beyond the view of the treeline as a single elevational boundary to jointly characterize forest infilling and densification process in multiple ecotone boundaries across the Pyrenees. Delpouve et al. (2025) reported that land abandonment strongly influenced the velocity of forestline shifts, with the closed forest advancing faster than the forest line, a pattern that closely parallels our own observation of faster timberline than tree species line advance. Together with Hooshyarkhah et al. (2025), these studies indicate that treeline ecotones are best understood not as single lines but as heterogeneous zones whose internal structure, and not only their outer boundary, is reorganising in response to land-use and climatic change.

### 4.2 | Land use as the primary driver of vegetation dynamics in Alpine treelines

Whether climate warming or land-use change act as the primary drivers of recent treeline advance remains an open and actively debated question in treeline ecology (Nicoud et al., 2025; Marzini et al., 2025). The long-term, multi-site analysis of timberline, tree species line, and ecotonal surface dynamics here employed tests their relative contribution, and provides evidence that historical land use plays a major role in shaping treeline dynamics in the Italian Alps.

The faster timberline shift is consistent with a long history of traditional practices that depressed timberlines below their climatic potential (Sanjuán et al., 2018). The widespread abandonment since the mid-twentieth century (Anselmetto et al., 2024; Lasanta et al., 2017) released the ecotone’s lower boundary from anthropogenic pressure, promoting rapid recolonization of former alpine grasslands (Anselmetto et al., 2022; Tasser et al., 2007). By contrast, tree species lines likely experienced weaker historical pressure due to their greater remoteness and steeper terrain (Lavorel et al., 2017; Toivonen et al., 2011; Urquiaga-Flores et al., 2024), and their slower advance suggests they already sit closer to their climatic limit— consistent with the limited explanatory power of climatic variables for both boundaries. Patterns of ecotonal surface change further support this interpretation. The strongest ecotone contractions occurred in Senales, Devero, Livigno, and Becco (Fig. 6a), which also recorded the strongest timberline advance, whereas sites with stable or expanding ecotones showed smaller timberline shifts. These contrasting responses suggest that ecotone dynamics depend on the balance between the two boundaries and likely reflect differences in historical land-use intensity, timing of abandonment, and proximity to climatic limits.

The mean elevation of the 1954 timberline was the strongest predictor across all response variables, explaining one-half to two-thirds of observed variance (Fig. 7). Historically depressed timberlines had greater potential for upslope recovery once anthropogenic pressure declined, whereas sites with higher initial treelines were already closer to their climatic limit and shifted less. The negative relationship between baseline timberline elevation and elevational shift, together with its positive association with ecotonal surface contraction, indicates that present-day dynamics largely reflect the legacy of past land use. Total forest cover change points in the same direction, with landscapes undergoing greater afforestation also showing larger boundary advances and stronger ecotone contraction. Road density, used as a proxy for accessibility and persistence of human activity (Anselmetto et al., 2024; Toivonen et al., 2011), was negatively associated with elevational shift, suggesting slower recovery where disturbance persisted; human population density, previously identified as an indicator of forest expansion (Anselmetto et al., 2024; Delpouve et al., 2025), showed no significant effect here.

**Figure 7.**
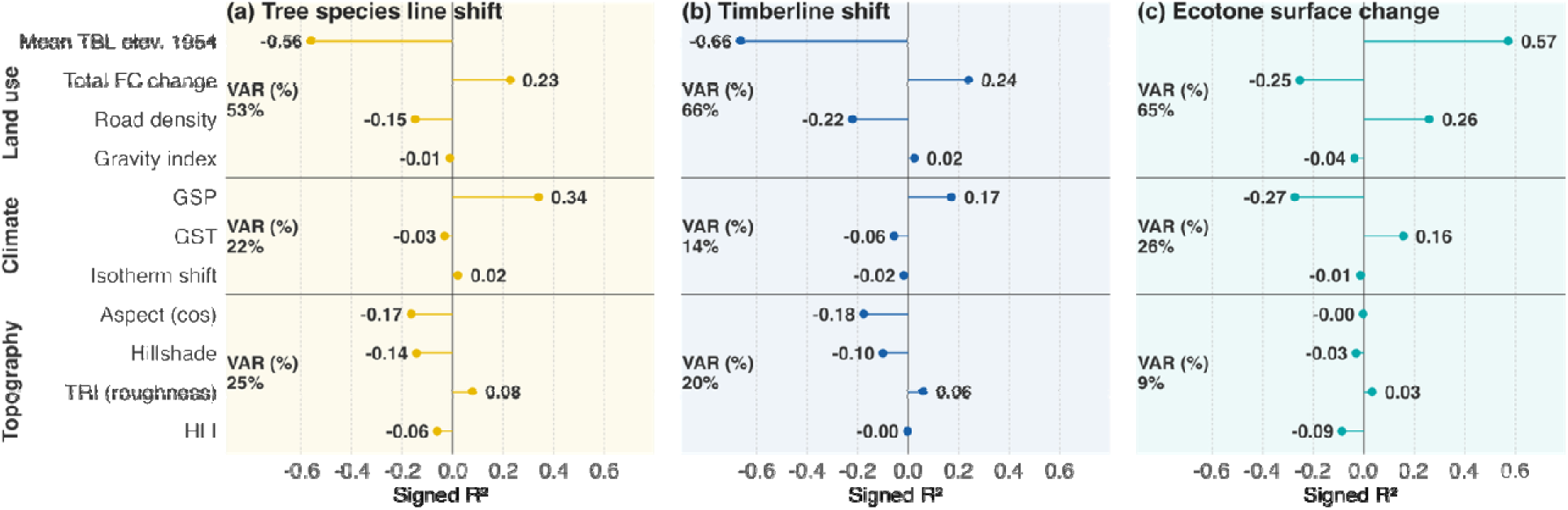
| Signed R² values from simple ordinary least squares (OLS) regressions relating each predictor to the mean per-transect elevational shift of the tree species line (a), the timberline (b), and ecotone surface change (c), across nine study sites. The length of each lollypop stem indicates the proportion of variance explained (R²), and the sign reflects the direction of the OLS slope (positive values indicate that higher predictor values are associated with greater upslope shift or surface expansion). All predictors were normalised prior to modelling to allow direct comparison of effect sizes across variables measured on different scales. GST = mean growing-season temperature; GSP = growing-season precipitation; TRI = topographic roughness index; HLI = heat load index; Mean TBL elev. 1954 = mean elevation of the 1954 timberline; Total FC change = proportional change in dense forest cover between 1954 and 2017.

Climatic variables explained substantially less variance than land-use variables, but were not without effect. GSP was the strongest climatic predictor, consistent with its role in forest regeneration and seedling establishment (Dirnböck et al., 2003; He et al., 2023; Xie et al., 2024), whereas growing-season temperature and isotherm shift had limited explanatory power despite well-documented Alpine warming (Beniston et al., 1997). GSP contributed more to tree species line than to timberline shifts (Fig. 7a,b), suggesting that timberline displacement is governed primarily by land-use legacy, whereas the tree species line— historically less human-constrained and located in harsher, more climatically limited terrain—is comparatively more sensitive to water availability. GSP also showed a marked negative contribution to ecotonal surface change, indicating that wetter conditions are associated with greater contraction. Wetter conditions might foster timberline vegetation by ameliorating growing conditions, accelerating its upward shift in a greater way than the tree species line. Overall, in our study sites climate conditions appear to modulate treeline change within a trajectory primarily set by land abandonment, rather than acting as an independent driver of comparable strength.

### 4.3 | Limitations and conclusive remarks

This study provides evidence that in historically intensively managed Alpine landscapes, vegetation and treeline dynamics still respond to past land use and its relaxation rather than recent climate warming. Treelines not only recorded an upward displacement of both its boundaries but also a measurable contraction of the ecotonal surface. The predominance of disturbance-related predictors and the limited role of temperature metrics are consistent with our hypothesis of secondary succession following the cessation of human disturbance, rather than a climate-driven process. Generalising this conclusion requires caution, especially in regions without a comparable strong land-use legacy.

Climate and land-use effects remain difficult to disentangle because warming and forest expansion occurred concurrently during the study period.

Treeline ecotone dynamics were assessed based on a dataset of detectable trees in 1-meter resolution aerial imagery. Seedlings and saplings were therefore not included in the analyses, but the comparison of historical and recent aerial images with the same spatial resolution makes the detected dynamics reliable.

Future research should combine broader spatial coverage with high-resolution land-use and climate data to better resolve the relative importance of these drivers and account for local environmental heterogeneity.

The observed contraction of treeline ecotones may have important consequences for biodiversity and ecosystem functioning in high-elevation grasslands, where many species depend on open, low-competition habitats. Continued land abandonment and warming may accelerate habitat loss and increase the risk of local extinctions, underscoring the value of monitoring ecotone surface dynamics, and not only boundary position, in future assessments of Alpine treeline change.

## ACKNOWLEDGMENTS

We sincerely thank Simone Ferrero for developing the land-cover maps during its master thesis. This research has been supported by the Ministero dell’Università e della Ricerca through the project OLYMPUS-Spatio-temporal analysis of Mediterranean treeline patterns: a multiscale approach (PRIN: PROGETTI DI RICERCA DI RILEVANTE INTERESSE NAZIONALE – Bando 2022 – project code: 20225S47P8).

## COMPETING INTERESTS

The authors declare no conflicts of interest.

## AUTHOR CONTRIBUTIONS

Erik Carrieri, Nicolò Anselmetto, Donato Morresi, and Matteo Garbarino conceived the ideas and designed the methodology; Fabio Meloni and Erik Carrieri collected and curated the data; Erik Carrieri, HaTrang Nguyen, Nicolò Anselmetto, Donato Morresi analysed and interpreted the data; Erik Carrieri led the writing of the manuscript; Matteo Garbarino led the OLYMPUS PRIN project. All authors contributed critically to the drafts and gave final approval for publication.

## DATA AVAILABILITY

The data that support the findings of this study are openly available in the Dryad Digital Repository: https://datadryad.org/share/LINK_NOT_FOR_PUBLICATION/TBVvXqHCSPGq_pYsRIMILTtVD_WuJHJGsoYruykwVSA

## Notes

https://doi.org/10.5061/dryad.8w9ghx43c

### Competing Interest Statement

The authors have declared no competing interest.

