## Supplementary materials for "Treeline ecotone habitat loss and upward shift is driven by land abandonment rather than climate change in the Italian Alps"

### Supplementary Material A – Supporting Information on study sites and aerial photographs

**Table S1 |** *Upper and lower limit, and total surface of the region of interest (ROI) in each study site. The ROI corresponds to the region within which the tree species line, the timberline, and the treeline ecotone delineation was performed.*

| Site | Lower boundary | Upper boundary | ROI surface (ha) |
| --- | --- | --- | --- |
| becco | 2000 | 2500 | 411.9 |
| bocche | 2000 | 2400 | 475.8 |
| chianale | 2000 | 2500 | 278.9 |
| devero | 2000 | 2500 | 577 |
| genevris | 2000 | 2500 | 441.7 |
| livigno | 2000 | 2500 | 377.8 |
| rion | 2000 | 2500 | 228.3 |
| senales | 2000 | 2500 | 444.8 |
| valfurva | 2000 | 2500 | 321.6 |
| Italian Alps | **/** | **/** | **3557.8** |

**Aerial photographs acquisition and processing.** Grayscale aerial photographs acquired during the 1954 national flight campaign (GAI flight) commissioned by the Italian Geographic Military Institute (IGM) and RGB orthophotos acquired in 2017 from Agenzia per le Erogazioni in Agricoltura (AGEA) were used to detect treeline shifts. Aerial photographs acquired in 1954 were captured from an altitude of ≈10,000 m (equivalent to a scale of 1:53,000) using 23 × 23 cm film frames and presented a pixel size of 1-meter. Recent RGB orthophotos were acquired with high-resolution digital cameras and feature a pixel size between 20 and 50 cm. All aerial photographs were scanned at 2400 DPI, georeferenced and orthorectified in PCI Geomatica 10.2 using a 10-m resolution DEM (TINItaly DEM; Tarquini et al., 2007). All images were projected to WGS84/UTM Zone 32N (EPSG:32632). To ensure comparability between datasets, all imagery was resampled to 1-m spatial resolution. Within each study site, we defined regions of interest (ROIs) as areas between 2000 and 2500 m a.s.l., an elevation range widely documented as the transition zone from closed forest to the uppermost isolated trees in the European Alps (Körner and Hiltbrunner, 2024; Malfasi and Cannone, 2020; Paulsen et al., 2000).

### Supplementary Material B – Supporting Information on treeline detection

**Treeline Detection.** To delineate the timberline and tree species line we produced a spatially explicit representation of tree crowns by classifying tree and non-tree pixels in both the 1954 and 2017 aerial photographs. The workflow for the production of the binary tree cover layer for each time period and site consisted in: (i) manual annotation of training samples, (ii) training of the deep learning model, and (iii) post-processing of models predictions. The binary layers resulting from these steps formed the foundation data from which the timberline and the tree species line were delineated.

**Training and validation dataset.** Training and validation samples were generated by overlaying a 256 × 256 m grid on each orthophoto and selecting tiles Intersecting the ROI (Figure 2). The 256-m tile size balanced capture of fine-scale crown morphology with computational efficiency during deep-learning inference. To minimize spatial autocorrelation and redundant information, adjacent tiles were excluded. Sampling was stratified to represent variability in crown morphology, illumination, and background texture across the study area. Tile selection aimed to encompass the full range of tree spectral information in the orthophotos because annotation quality strongly affects convolutional-neural-network performance (Fassnacht et al., 2016; Nguyen et al., 2024). For historical grayscale imagery, tiles were chosen to represent the full gray tonal range of tree crowns, whereas for recent RGB imagery, selection prioritized variation in vegetation colour, illumination, and shadow patterns known to influence segmentation accuracy (Nguyen et al., 2024).

A total of 96 RGB tiles and 91 grayscale tiles were manually annotated into a binary tree/background mask using Anylabelling (Kirillov et al., 2023). In this study we defined individual tree crowns as rounded or irregularly circular features exhibiting green–brown tonal patterns in RGB imagery or medium-gray reflectance in the 1954 photographs. In high-elevation environments, sun angle and canopy geometry often produce characteristic shadow signatures (Figure S1). The elongated, conical shadows of isolated crowns were used as an additional cue to aid annotation (Fassnacht et al., 2016; Lin et al., 2024). Rocky surfaces tend to appear angular or polygonal and cast shorter, less directional shadows, which together with the geometric and tonal cues, helped reduce misclassification between tree crowns and exposed rock (Nguyen et al., 2024). Because the objective was to produce spatially consistent individual tree crown predictions for ecotone analyses rather than to benchmark model architectures, the annotated dataset was partitioned into 80% for training, 10% for validation and 10% for testing.

### **Deep learning model.** We applied a deep learning approach based on a convolutional neural network (CNN) trained on historical and recent aerial imagery to assess changes in tree cover over time. Binary land-cover classification and ecotonal tree-cover mapping were performed with a UNet++ deep convolutional neural network (Zhou et al., 2018). Two distinct models were trained, one per year. Grayscale images were stacked into three identical channels prior to training so that both models shared the same internal architecture. The encoder was an SE-ResNeXt101 pretrained on ImageNet (Xie et al., 2017), which extracted multiscale features; the decoder was an scSE-enhanced UNet++ (Sahragard et al., 2025) that reconstructed detailed spatial patterns. Models were optimized using the Lovász loss to maximize Intersection-over-Union with the Adam optimizer at a learning rate of 1×10⁻⁵. Performance was evaluated with IoU, F1 score, and Cohen’s Kappa on the training and validation sets. For inference over large orthophotos, predictions were generated on overlapping 256 × 256 pixel tiles and merged using Gaussian-weighted blending to minimize edge artifacts.

### **Classification post-processing.** Model outputs were manually post-processed to correct residual classification errors. False positives—typically small, clustered patches beyond the treeline on bare ground or rocky slopes—were removed by manual correction (Figure S2, Table S1). Under-classification generally appeared as gaps within dense forest where overlapping shadows fragmented the canopy mask (Figure S3). Such illumination-related artifacts are common in high-resolution mountain imagery because low solar angles increase shadow extent and reduce the apparent fraction of sunlit vegetation (Nguyen et al., 2024). To homogenize dense stands and compensate for shadow-induced gaps, focal statistics were applied. Pixels with a local canopy cover density ≥30% within a 10 m radius were reclassified as “tree,” thereby filling shadow-affected voids and restoring a more realistic dense-forest representation (Figure S2).

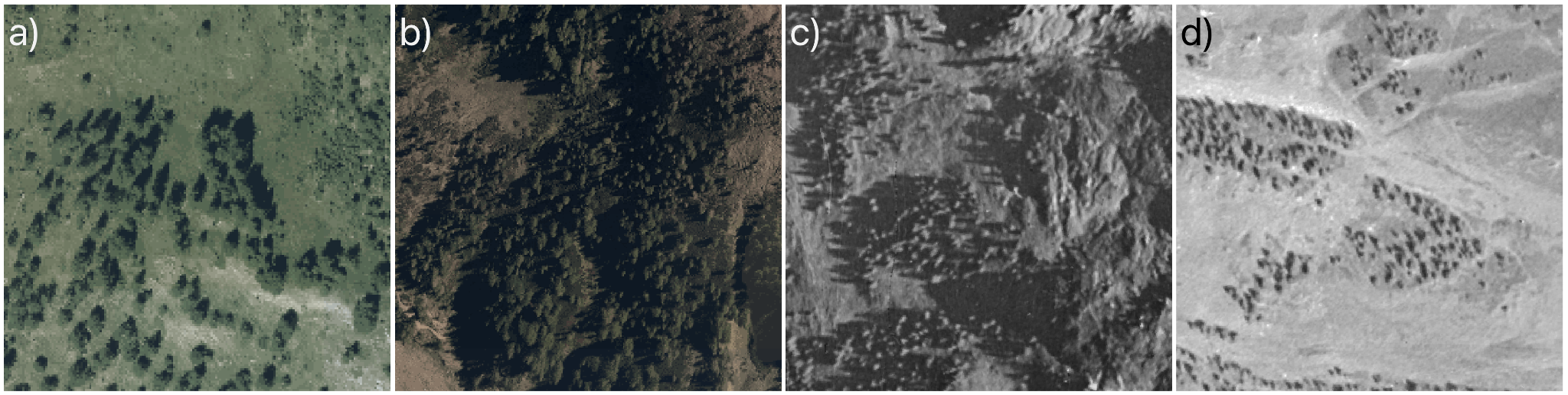
**Figure S1** | *Variability of shades of tree crown colors and shadow length in (a,b) RGB images and (c,d) greyscale images under different lighting conditions, camera angle, and spectral response.*

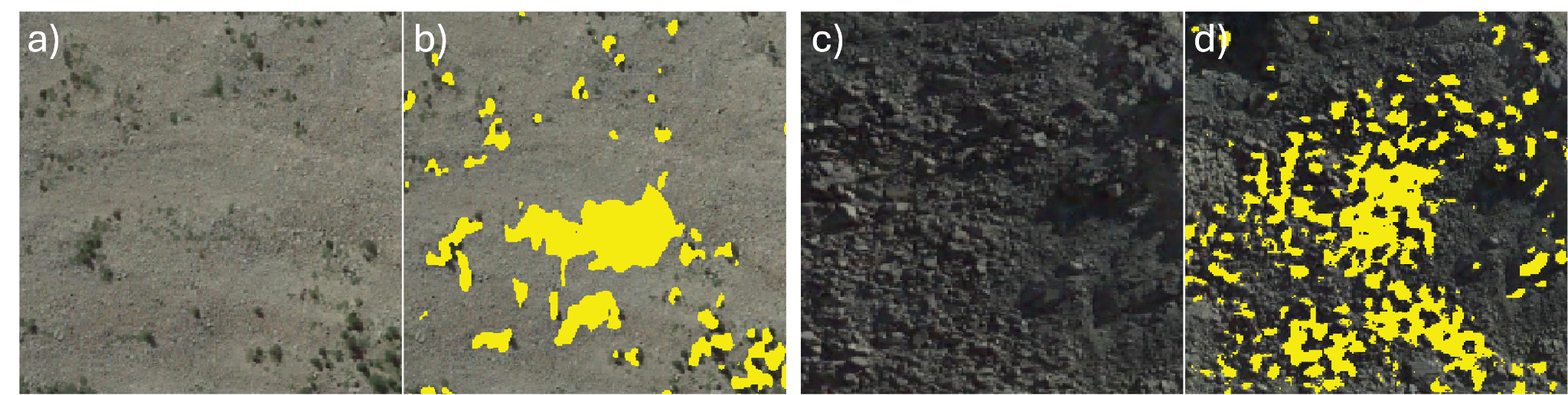
**Figure S2** | *Misclassification examples. (a) bare ground and (c) rocks misinterpreted as trees (b) and (d).*

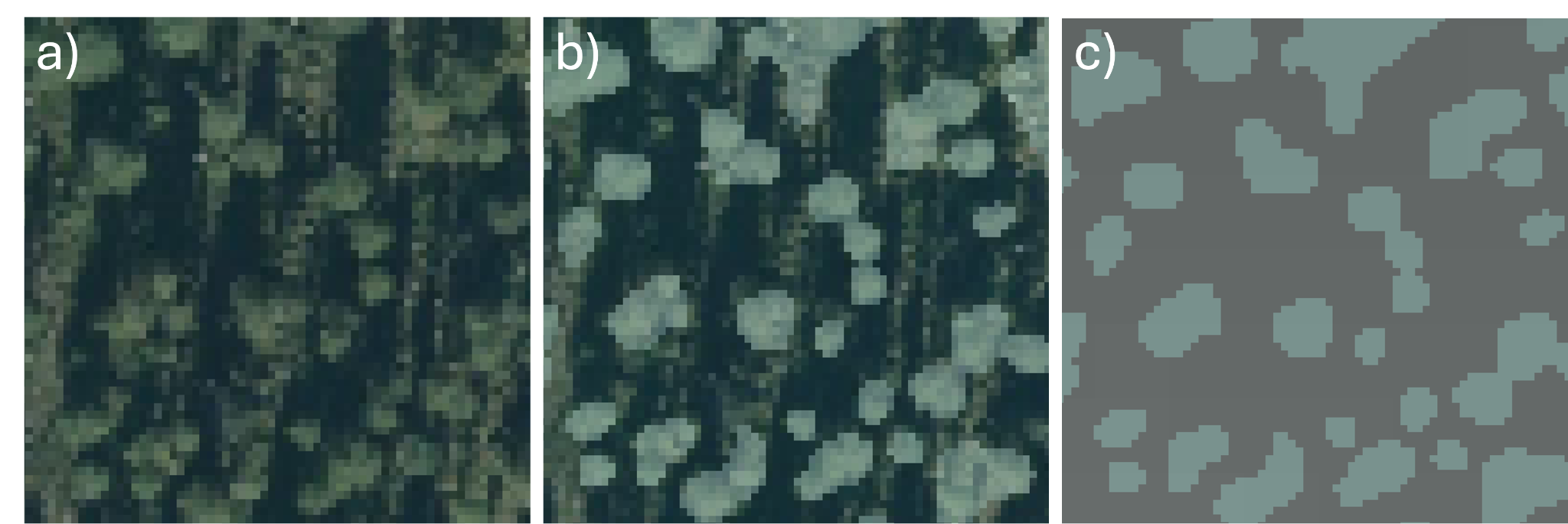

**Figure S3** *| (a) dense forest with shadow gaps in RGB images, (b) classification results of tree detection overlaid with the RGB image, and (c) dense forest patches classifications appearance*.

**Table S2** | *Classification results for tree detection. We report Intersection over Union (IoU), F1-score, User Accuracy (UA) and Cohen's Kappa for classification of trees on RGB and greyscale images.*

| **Year** | **Class name** | **IoU** | **F1** | **UA** | **κ** |
| --- | --- | --- | --- | --- | --- |
| 1954 | Background | 0.81 | 0.89 | 0.88 | - |
|  | Trees | 0.50 | 0.66 | 0.70 | - |
|  | Weighted average | 0.73 | 0.83 | 0.83 | 0.56 |
| 2017 | background | 0.91 | 0.95 | 0.95 | - |
|  | trees | 0.54 | 0.70 | 0.72 | - |
|  | Weighted average | 0.86 | 0.92 | 0.92 | 0.65 |

**APPENDIX A** **|** References

Fassnacht, F. E., Latifi, H., Stereńczak, K., Modzelewska, A., Lefsky, M., Waser, L. T., Straub, C., and Ghosh, A.: Review of studies on tree species classification from remotely sensed data, Remote Sensing of Environment, 186, 64–87, https://doi.org/10.1016/j.rse.2016.08.013, 2016.

Hoover, E. M.: An introduction to regional economics, New York : Knopf, 472 pp., 1984.

Kirillov, A., Mintun, E., Ravi, N., Mao, H., Rolland, C., Gustafson, L., Xiao, T., Whitehead, S., Berg, A. C., Lo, W.-Y., Dollár, P., and Girshick, R.: Segment Anything, in: 2023 IEEE/CVF International Conference on Computer Vision (ICCV), 2023 IEEE/CVF International Conference on Computer Vision (ICCV), 3992–4003, https://doi.org/10.1109/ICCV51070.2023.00371, 2023.

Kline, J. D., Moses, A., and Alig, R. J.: Integrating Urbanization into Landscape-level Ecological Assessments, Ecosystems, 4, 3–18, https://doi.org/10.1007/s100210000056, 2001.

Lin, Y., Li, H., Jing, L., Ding, H., and Tian, S.: Individual Tree Crown Delineation Using Airborne LiDAR Data and Aerial Imagery in the Taiga–Tundra Ecotone, Remote Sensing, 16, 3920, https://doi.org/10.3390/rs16213920, 2024.

Nguyen, T.-A., Rußwurm, M., Lenczner, G., and Tuia, D.: Multi-temporal forest monitoring in the Swiss Alps with knowledge-guided deep learning, Remote Sensing of Environment, 305, 114109, https://doi.org/10.1016/j.rse.2024.114109, 2024.

Polyakov, M., Majumdar, I., and Teeter, L.: Spatial and temporal analysis of the anthropogenic effects on local diversity of forest trees, Forest Ecology and Management, 255, 1379–1387, https://doi.org/10.1016/j.foreco.2007.10.052, 2008.

Sahragard, E., Farsi, H., and Mohamadzadeh, S.: Advancing semantic segmentation: Enhanced UNet algorithm with attention mechanism and deformable convolution, PLoS One, 20, e0305561, https://doi.org/10.1371/journal.pone.0305561, 2025.

Xie, S., Girshick, R., Dollar, P., Tu, Z., and He, K.: Aggregated Residual Transformations for Deep Neural Networks, in: 2017 IEEE Conference on Computer Vision and Pattern Recognition (CVPR), 2017 IEEE Conference on Computer Vision and Pattern Recognition (CVPR), 5987–5995, https://doi.org/10.1109/CVPR.2017.634, 2017.

Zhou, Z., Rahman Siddiquee, M. M., Tajbakhsh, N., and Liang, J.: Unet++: 4th International Workshop on Deep Learning in Medical Image Analysis, DLMIA 2018 and 8th International Workshop on Multimodal Learning for Clinical Decision Support, ML-CDS 2018 Held in Conjunction with MICCAI 2018, Deep Learning in Medical Image Analysis and Multimodal Learning for Clinical Decision Support - 4th International Workshop, DLMIA 2018 and 8th International Workshop, ML-CDS 2018 Held in Conjunction with MICCAI 2018, 3–11, https://doi.org/10.1007/978-3-030-00889-5_1, 2018.

### Supplementary Material C – Supporting Information on materials and methods

The final suite of predictors (Table 1) was defined after the computation of pairwise Pearson correlations across all candidate variables. Variables with r > 0.80 with a retained variable were excluded.

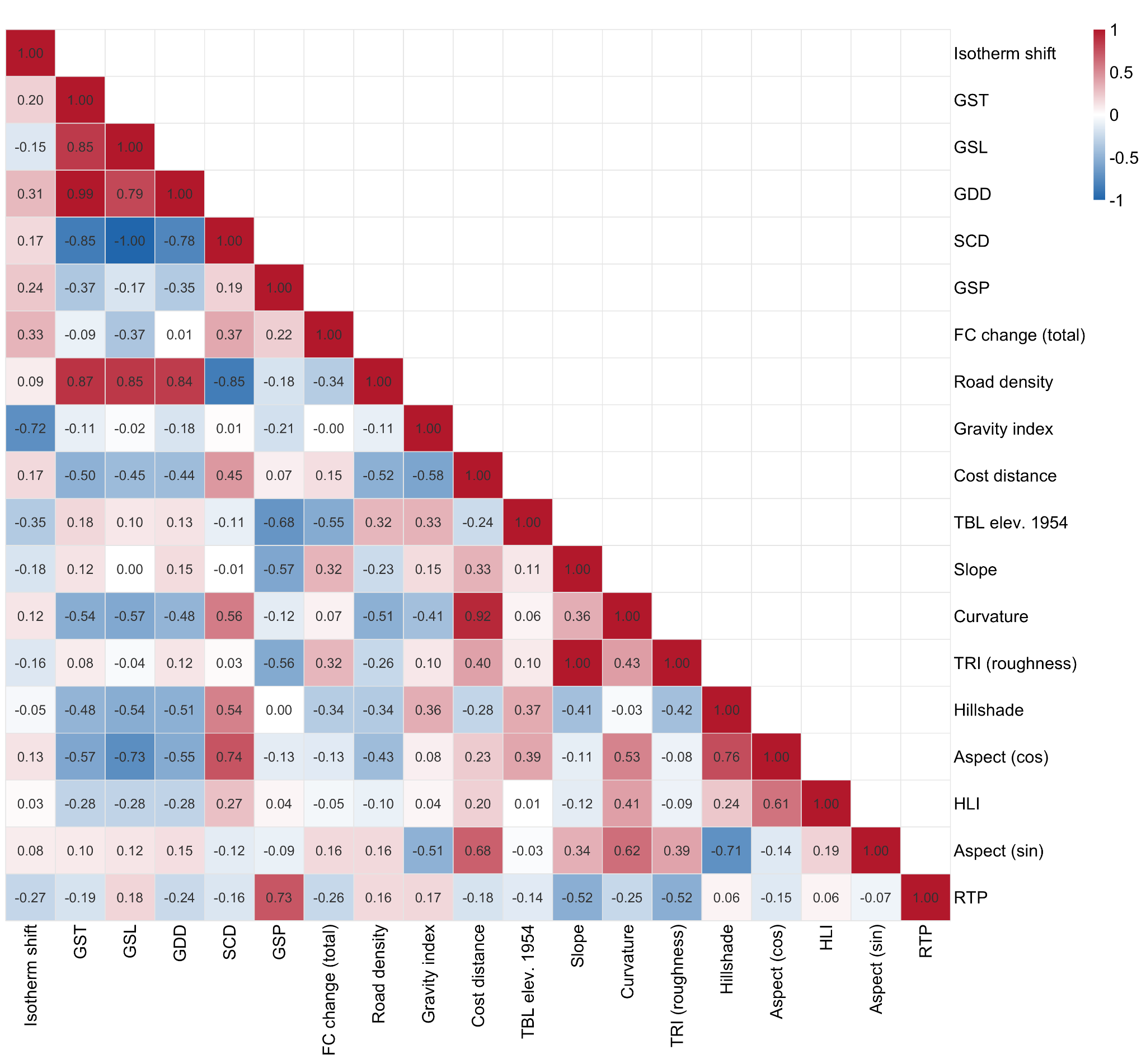

**Figure S4 |** *Pearson correlation matrix among all the 19 predictor variables. Correlations were computed using pairwise complete observations (n = 9 sites). Color scale ranges from strong negative (blue, r = −1) to strong positive (red, r = 1). Acronyms: growing-season temperature (GST); growing season length (GSL); growing degree days (GDD); snow cover duration (SCD); growing-season precipitation (GSP); forest cover (FC); timberline (TBL); topographic ruggedness index (TRI); heat load index (HLI); relative topographic position (RTP).*

**Population gravity index (gravity index).** The population gravity index implemented among the land-use predictors accounts for the presence and the size of nearby cities as well as their proximity to the treelines. In this study the gravity index is considered as a proxy of human pressure on the treeline area, as more densely populated areas are often associated with higher intensity and frequency of traditional activities —e.g. grazing, fire use, wood cutting, mowing— in surrounding rural areas (Anselmetto et al., 2024; Wang et al., 2022). It is highest when large human populations are located nearby and decreases when populations are smaller or located further away.

We used traditional specification of population gravity index (PGI) suggested by Hoover (1984)

| ${Gravity index}_{i}\sum_{k} \frac{P_{k}}{D_{ki}^{2}}$ |
| --- |

where the gravity index for the point *i,* P*k* is the population of populated place *k*, and D*ki* is the distance between point *i* and populated place *k*. The data on population and distance from settlements was derived from OSM (Table S2). This index has been used by several studies to investigate the effect of human pressure and land-use change on forest ecosystems (Kline et al., 2001; Polyakov et al., 2008).

**Table S3** | *Built- up areas and road types and description from OpenStreetMap (OSM).*

| **Class** | **Name** | **Description** |
| --- | --- | --- |
| Built-up areas | City | The largest urban settlement or settlements within the territory. |
|  | Town | An important urban centre, between a village and a city in size. |
|  | Village | A smaller distinct settlement, smaller than a town with few facilities available with people traveling to nearby towns to access these. |
|  | Hamlet | A smaller rural community, typically with fewer than 100-200 inhabitants, and little infrastructure |
|  | Isolated dwelling | The smallest kind of settlement (1-2 households) |
|  | Farm | An individually named farm |
| Roads | Motorway | A restricted access major divided highway, normally with 2 or more running lanes plus emergency hard shoulder. Equivalent to the Freeway, Autobahn, etc.. |
|  | Trunk | The most important roads in a country's system that aren't motorways. (Need not necessarily be a divided highway.) |
|  | Primary | The next most important roads in a country's system. (Often link larger towns.) |
|  | Secondary | The next most important roads in a country's system. (Often link towns.) |
|  | Tertiary | The next most important roads in a country's system. (Often link smaller towns and villages) |
|  | Unclassified | The least important through roads in a country's system – i.e. minor roads of a lower classification than tertiary, but which serve a purpose other than access to properties. (Often link villages and hamlets.) |
|  | Track | Roads for mostly agricultural or forestry uses. |

### **APPENDIX C** **|** References

Anselmetto, N., Weisberg, P. J., and Garbarino, M.: Global change in the European Alps: A century of post-abandonment natural reforestation at the landscape scale, Landscape and Urban Planning, 243, 104973, https://doi.org/10.1016/j.landurbplan.2023.104973, 2024b.

Hoover, E. M.: An introduction to regional economics, New York : Knopf, 472 pp., 1984.

Kline, J. D., Moses, A., and Alig, R. J.: Integrating Urbanization into Landscape-level Ecological Assessments, Ecosystems, 4, 3–18, https://doi.org/10.1007/s100210000056, 2001.

Polyakov, M., Majumdar, I., and Teeter, L.: Spatial and temporal analysis of the anthropogenic effects on local diversity of forest trees, Forest Ecology and Management, 255, 1379–1387, https://doi.org/10.1016/j.foreco.2007.10.052, 2008.

Wang, X., Tao, W., Xu, J., Shen, Z., Yang, Y., Chen, A., Wang, S., Liang, E., and Piao, S.: Enhanced habitat loss of the Himalayan endemic flora driven by warming-forced upslope tree expansion, Nature Ecology & Evolution, 6, 1–10, https://doi.org/10.1038/s41559-022-01774-3, 2022.

### Supplementary Material D – Supporting Information on Results

**Table S4 |** *Results of residual diagnostic tests applied to each simple OLS regression model. For each predictor–response combination we report the Shapiro-Wilk statistic (SW) and associated p-value, and the Breusch-Pagan statistic (BP) and associated p-value for heteroscedasticity of residuals. Values in bold indicate p < 0.05.*

| **Driver type** | **Predictor** | **Response** | **SW** | **SW p** | **BP** | **BP p** |
| --- | --- | --- | --- | --- | --- | --- |
| Climate | GSP | **Timberline shift** | **0.675** | **0.001** | **0.32** | **0.572** |
|  |  | Tree species line shift | 0.842 | 0.061 | 0.208 | 0.649 |
|  |  | Ecotone surface change | 0.89 | 0.202 | 0.861 | 0.353 |
|  | GST | **Timberline shift** | **0.713** | **0.002** | **0.09** | **0.764** |
|  |  | Tree species line shift | 0.944 | 0.623 | 0.024 | 0.878 |
|  |  | Ecotone surface change | 0.979 | 0.956 | 1.688 | 0.194 |
|  | Isotherm shift | **Timberline shift** | **0.742** | **0.004** | **0.162** | **0.688** |
|  |  | Tree species line shift | 0.957 | 0.763 | 0.461 | 0.497 |
|  |  | Ecotone surface change | 0.913 | 0.338 | 0.123 | 0.725 |
| Land use | Cost of Movement | **Timberline shift** | **0.8** | **0.021** | **0.952** | **0.329** |
|  |  | Tree species line shift | 0.927 | 0.451 | 0.295 | 0.587 |
|  |  | Ecotone surface change | 0.88 | 0.157 | 0.429 | 0.512 |
|  | Mean TBL elev. 1954 | Timberline shift | 0.902 | 0.267 | 5.334 | 0.021 |
|  |  | Tree species line shift | 0.928 | 0.465 | 0.963 | 0.326 |
|  |  | Ecotone surface change | 0.94 | 0.577 | 0.734 | 0.392 |
|  | Road density | **Timberline shift** | **0.792** | **0.017** | **2.291** | **0.13** |
|  |  | Tree species line shift | 0.932 | 0.502 | 1.831 | 0.176 |
|  |  | Ecotone surface change | 0.918 | 0.374 | 3.769 | 0.052 |
|  | Total FC change | Timberline shift | 0.852 | 0.078 | 2.156 | 0.142 |
|  |  | Tree species line shift | 0.961 | 0.803 | 0.69 | 0.406 |
|  |  | Ecotone surface change | 0.894 | 0.222 | 2.704 | 0.1 |
| Topography | HLI | **Timberline shift** | **0.743** | **0.004** | **0.008** | **0.93** |
|  |  | Tree species line shift | 0.943 | 0.614 | 0.059 | 0.808 |
|  |  | Ecotone surface change | 0.901 | 0.256 | 0.044 | 0.834 |
|  | Aspect (cos) | Timberline shift | 0.92 | 0.396 | 3.557 | 0.059 |
|  |  | Tree species line shift | 0.966 | 0.861 | 1.577 | 0.209 |
|  |  | Ecotone surface change | 0.911 | 0.325 | 0.125 | 0.724 |
|  | Hillshade | Timberline shift | 0.911 | 0.323 | 3.911 | 0.048 |
|  |  | Tree species line shift | 0.917 | 0.366 | 0.956 | 0.328 |
|  |  | Ecotone surface change | 0.933 | 0.515 | 0.636 | 0.425 |
|  | TRI (roughness) | Timberline shift | 0.908 | 0.303 | 4.166 | 0.041 |
|  |  | Tree species line shift | 0.954 | 0.729 | 1.951 | 0.162 |
|  |  | Ecotone surface change | 0.86 | 0.096 | 2.384 | 0.123 |

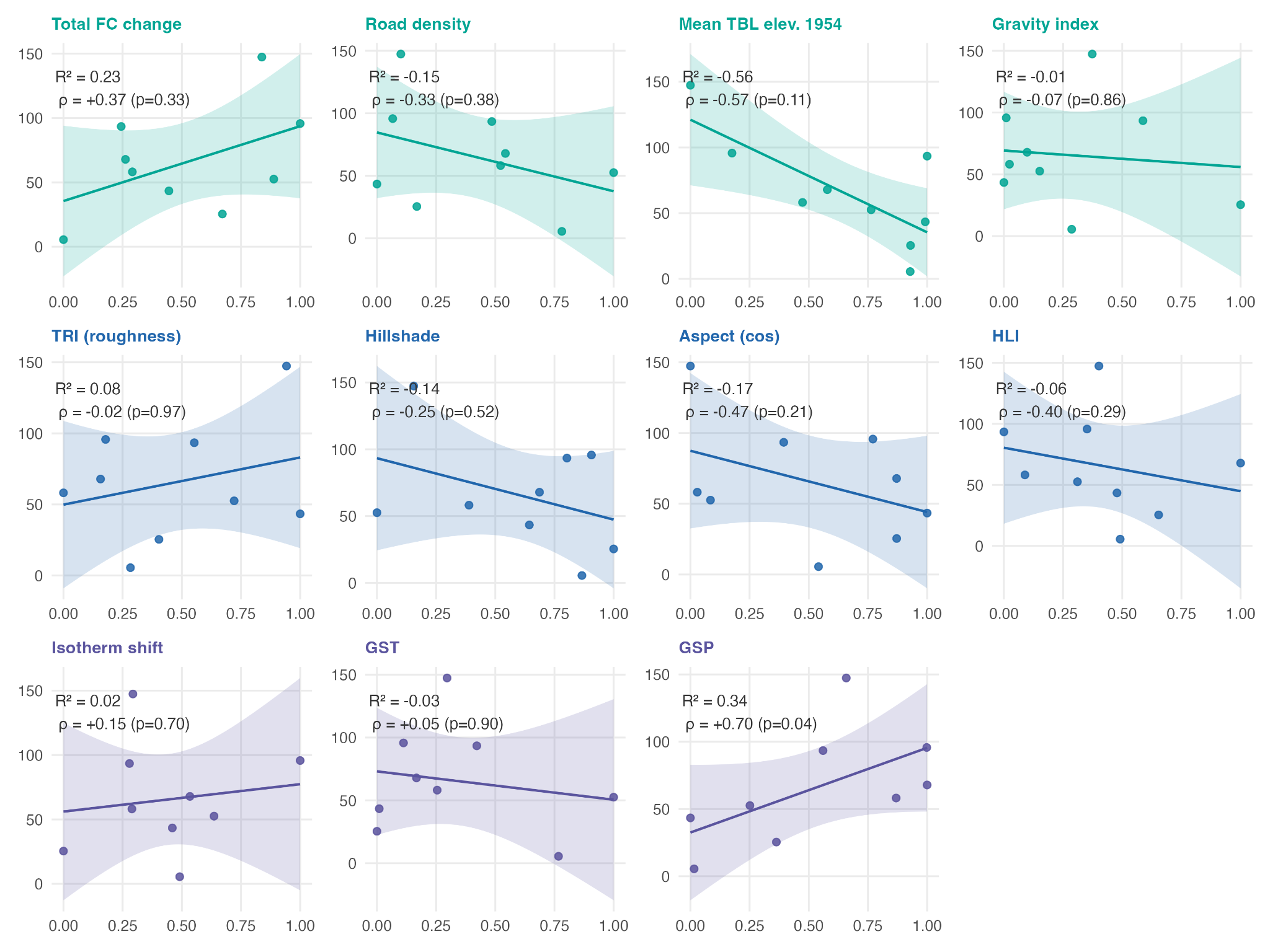

**Figure S5 |** *Bivariate relationships between each predictor and the tree species line elevational shift. Panels are grouped by driver type. Each point represents one of the nine study sites. The solid line shows the fitted ordinary least squares regression, and the shaded ribbon shows its 95% confidence interval. The R² value reported in each panel is signed according to the direction of the OLS slope; ρ denotes the Spearman rank correlation coefficient and its associated p-value.*

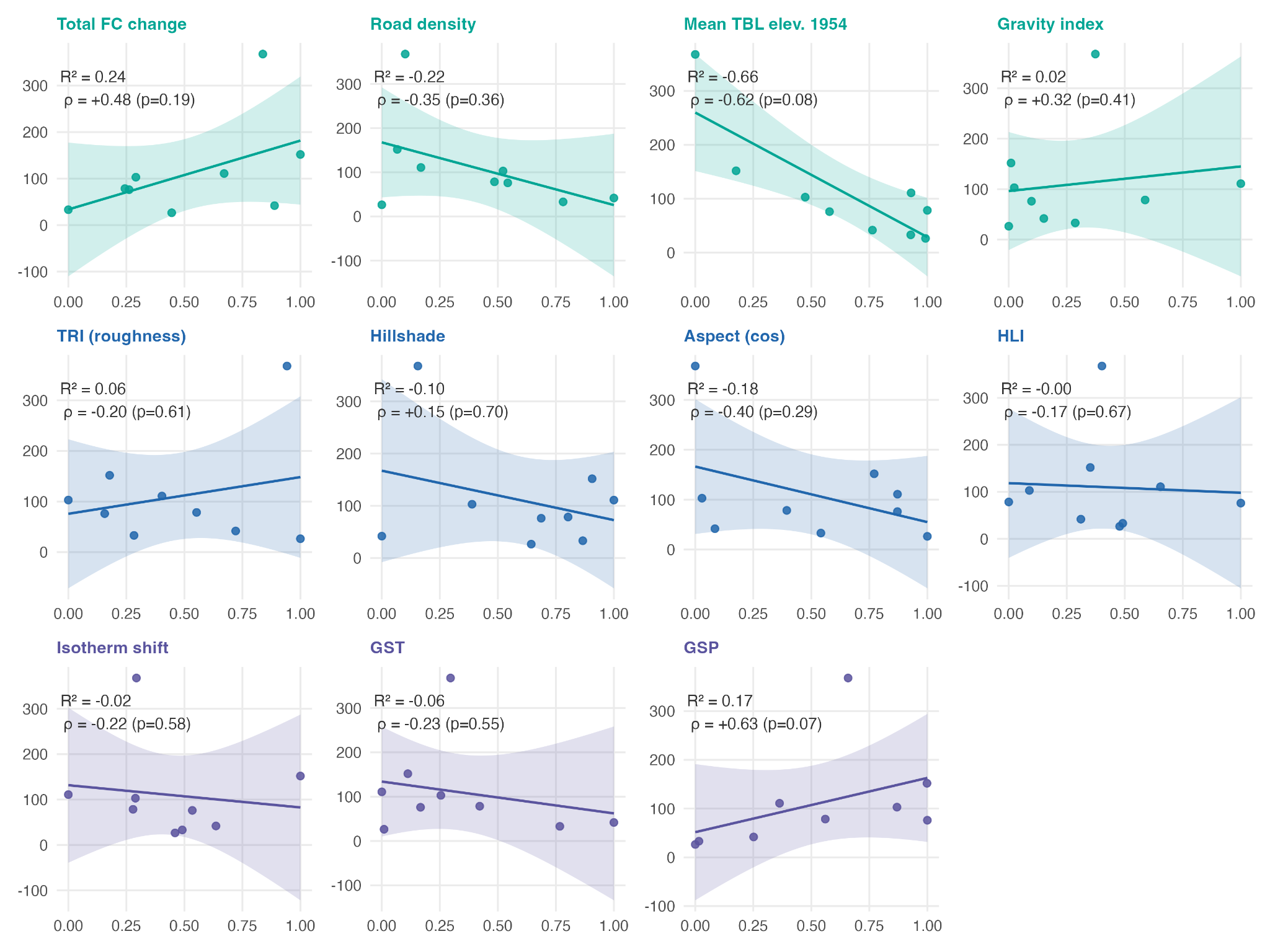

**Figure S6 |** *Bivariate relationships between each predictor and the timberline elevational shift. Panels are grouped by driver type. Each point represents one of the nine study sites. The solid line shows the fitted ordinary least squares regression, and the shaded ribbon shows its 95% confidence interval. The R² value reported in each panel is signed according to the direction of the OLS slope; ρ denotes the Spearman rank correlation coefficient and its associated p-value.*

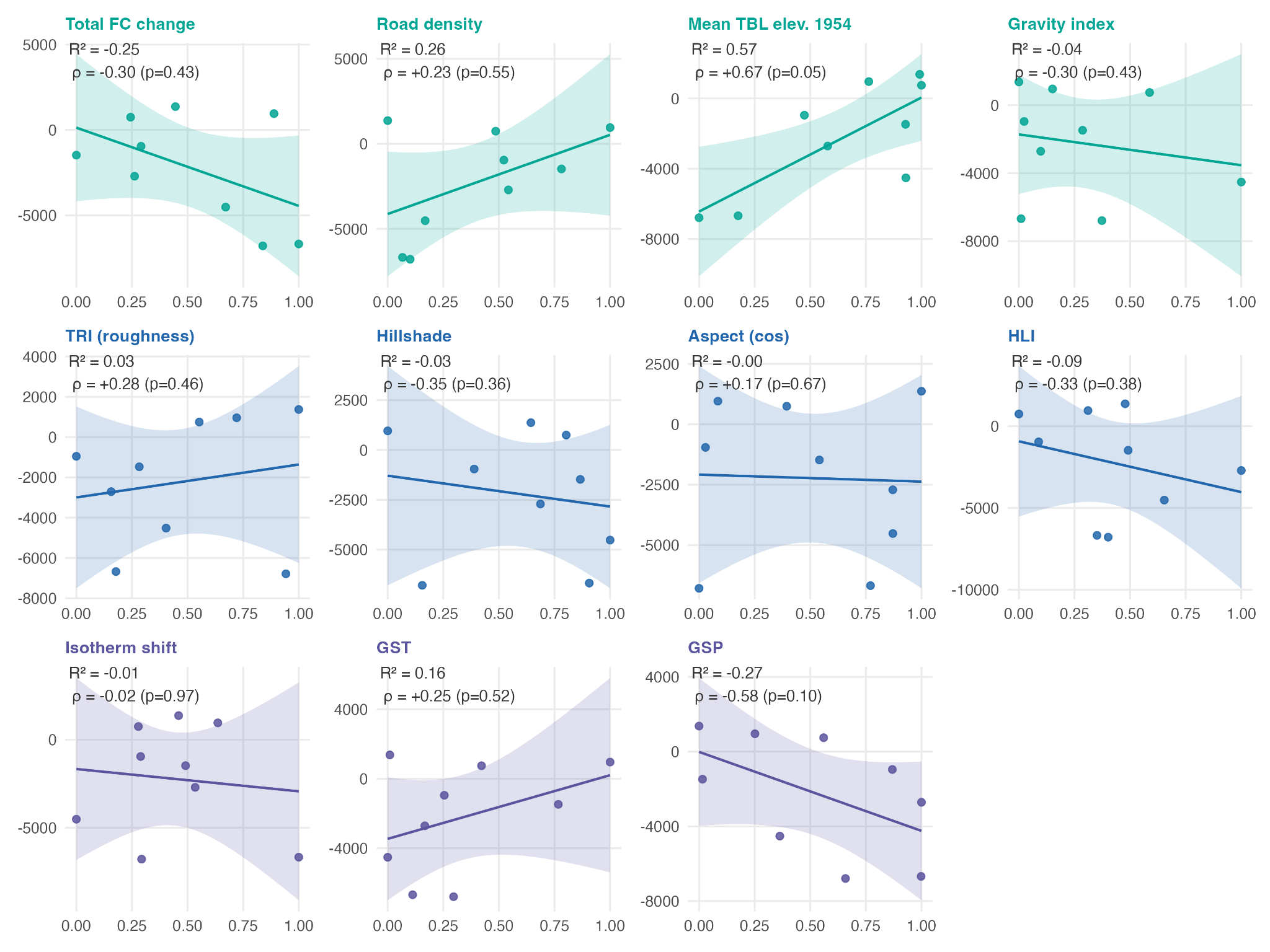

**Figure S7 |** B*ivariate relationships between each predictor and the ecotonal surface change. Panels are grouped by driver type. Each point represents one of the nine study sites. The solid line shows the fitted ordinary least squares regression, and the shaded ribbon shows its 95% confidence interval. The R² value reported in each panel is signed according to the direction of the OLS slope; ρ denotes the Spearman rank correlation coefficient and its associated p-value.*

**Table S5** **|** *Predictors explanatory power and direction of the effect on the three response variables. For each predictor we report the correlation coefficient R2, the OLS slope indicating the direction of the effect on the response variable, and spearman ρ and p indicating the strength of the association and statistical significance of the OLS result.*

| **Response** | **Predictor** | **R2** | **OLS slope** | **Spearman ρ** | | **Spearman p** |
| --- | --- | --- | --- | --- | --- | --- |
| Tree species line shift | Mean TBL elev. 1954 | 0.56 | -85.73 | -0.567 | 0.11 | |
|  | GSP | 0.34 | 62.94 | 0.7 | 0.04 | |
|  | Total FC change | 0.23 | 58.16 | 0.367 | 0.33 | |
|  | Aspect (cos) | 0.16 | -43.32 | -0.467 | 0.21 | |
|  | Road density | 0.15 | -47.06 | -0.333 | 0.38 | |
|  | Hillshade | 0.14 | -45.97 | -0.25 | 0.52 | |
|  | TRI (Roughness) | 0.08 | 33.19 | -0.017 | 0.97 | |
|  | HLI | 0.06 | -35.62 | -0.4 | 0.29 | |
|  | GST | 0.03 | -22.55 | 0.05 | 0.9 | |
|  | Isotherm shifts | 0.02 | 21.27 | 0.15 | 0.7 | |
|  | Gravity Index | 0.01 | -13.37 | -0.067 | 0.86 | |
| Timberline shift | Mean TBL elev. 1954 | 0.6 | -231.04 | -0.617 | 0.08 | |
|  | Total FC change | 0.24 | 148.15 | 0.483 | 0.19 | |
|  | Road density | 0.22 | -141.96 | -0.35 | 0.36 | |
|  | Aspect (cos) | 0.18 | -111.3 | -0.4 | 0.29 | |
|  | GSP | 0.17 | 111.27 | 0.633 | 0.07 | |
|  | Hillshade | 0.1 | -94.67 | 0.15 | 0.7 | |
|  | TRI (Roughness) | 0.06 | 72.53 | -0.2 | 0.61 | |
|  | GST | 0.06 | -71.81 | -0.233 | 0.55 | |
|  | Gravity Index | 0.02 | 48.8 | 0.317 | 0.41 | |
|  | Isotherm shifts | 0.02 | -49 | -0.217 | 0.58 | |
|  | HLI | 0 | -20.5 | -0.167 | 0.67 | |
| ESC | Mean TBL elev. 1954 | 0.57 | 6489.53 | 0.667 | 0.05 | |
|  | GSP | 0.27 | -4229.16 | -0.583 | 0.1 | |
|  | Road density | 0.26 | 4657.25 | 0.233 | 0.55 | |
|  | Total FC change | 0.25 | -4585.08 | -0.3 | 0.43 | |
|  | GST | 0.16 | 3665.22 | 0.25 | 0.52 | |
|  | HLI | 0.09 | -3107.17 | -0.333 | 0.38 | |
|  | Gravity Index | 0.04 | -1812.09 | -0.3 | 0.43 | |
|  | TRI (Roughness) | 0.03 | 1633.87 | 0.283 | 0.46 | |
|  | Hillshade | 0.03 | -1548.21 | -0.35 | 0.36 | |
|  | Isotherm shifts | 0.01 | -1269.22 | -0.017 | 0.97 | |
|  | Aspect (cos) | 0 | -288.56 | 0.167 | 0.67 | |
